# CaMKII binds both substrates and activators at the active site

**DOI:** 10.1101/2020.10.25.354241

**Authors:** Can Özden, Roman Sloutsky, Tomohiro Mitsugi, Nicholas Santos, Emily Agnello, Christl Gaubitz, Joshua Foster, Emily Lapinskas, Edward A. Esposito, Takeo Saneyoshi, Brian A. Kelch, Scott C. Garman, Yasunori Hayashi, Margaret M. Stratton

**Affiliations:** Department of Biochemistry and Molecular Biology, University of Massachusetts, Amherst, MA 01003, USA; Molecular and Cellular Biology Graduate Program, University of Massachusetts, Amherst, MA 01003, USA; Malvern Panalytical, Northampton, MA 01060; Department of Biochemistry and Molecular Biotechnology, University of Massachusetts Chan Medical School, Worcester, MA 01605, USA; Department of Pharmacology, Kyoto University Graduate School of Medicine, Kyoto, 606-8501 Japan

**Author notes:** Corresponding author. (M.M.S.).

**Keywords:** Ca^2+^/calmodulin dependent protein kinase II, X-ray crystallography, NMDA-type glutamate receptor, AMPA-type glutamate receptor, T-lymphoma invasion, metastasis-inducing protein 1

## Abstract

Ca^2+^/calmodulin dependent protein kinase II (CaMKII) is a signaling protein required for long-term memory. Once activated by Ca^2+^/CaM, it sustains activity even after the Ca^2+^ dissipates. In addition to well-known autophosphorylation-mediated mechanism, interaction with specific binding partners also persistently activates CaMKII. A longstanding model invokes two distinct S- and T-sites. If an interactor binds at the T-site, it will preclude autoinhibition and allow substrates to be phosphorylated at the S-site. Here, we specifically test this model with X-ray crystallography, molecular dynamics simulations, and biochemistry. Our data are inconsistent with this model. Co-crystal structures of four different activators or substrates show that they all bind to a single continuous site across the kinase domain. We propose a mechanistic model that persistent CaMKII activity is facilitated by high affinity binding partners, which kinetically compete with autoinhibition by the regulatory segment to allow substrate phosphorylation.

## INTRODUCTION

Ca^2+^/calmodulin-dependent protein kinase II (CaMKII) is a central signaling protein that controls cellular functions such as synaptic plasticity, cytoskeletal regulation, cell growth and division, gene transcription, and ion channel modulation [1]. CaMKII biology has been an active focus of research especially because of its crucial role in long-term potentiation (LTP), which is the basis for long-term memory [2, 3]. CaMKII is highly abundant in the forebrain postsynaptic density (PSD) fraction, where it makes up to 2% of the total protein [4]. CaMKII is a multisubunit complex made up of 12-14 subunits, which is oligomerized by the hub domain (**Fig. 1A**). Each CaMKII subunit contains a Ser/Thr kinase domain, autoinhibitory/regulatory segment, variable linker region, and a hub domain (**Fig. 1B**). CaMKII is governed by many molecular interactions, however, the structural details of these interactions are lacking (**Fig. 1C**) [5].

**Figure 1.**
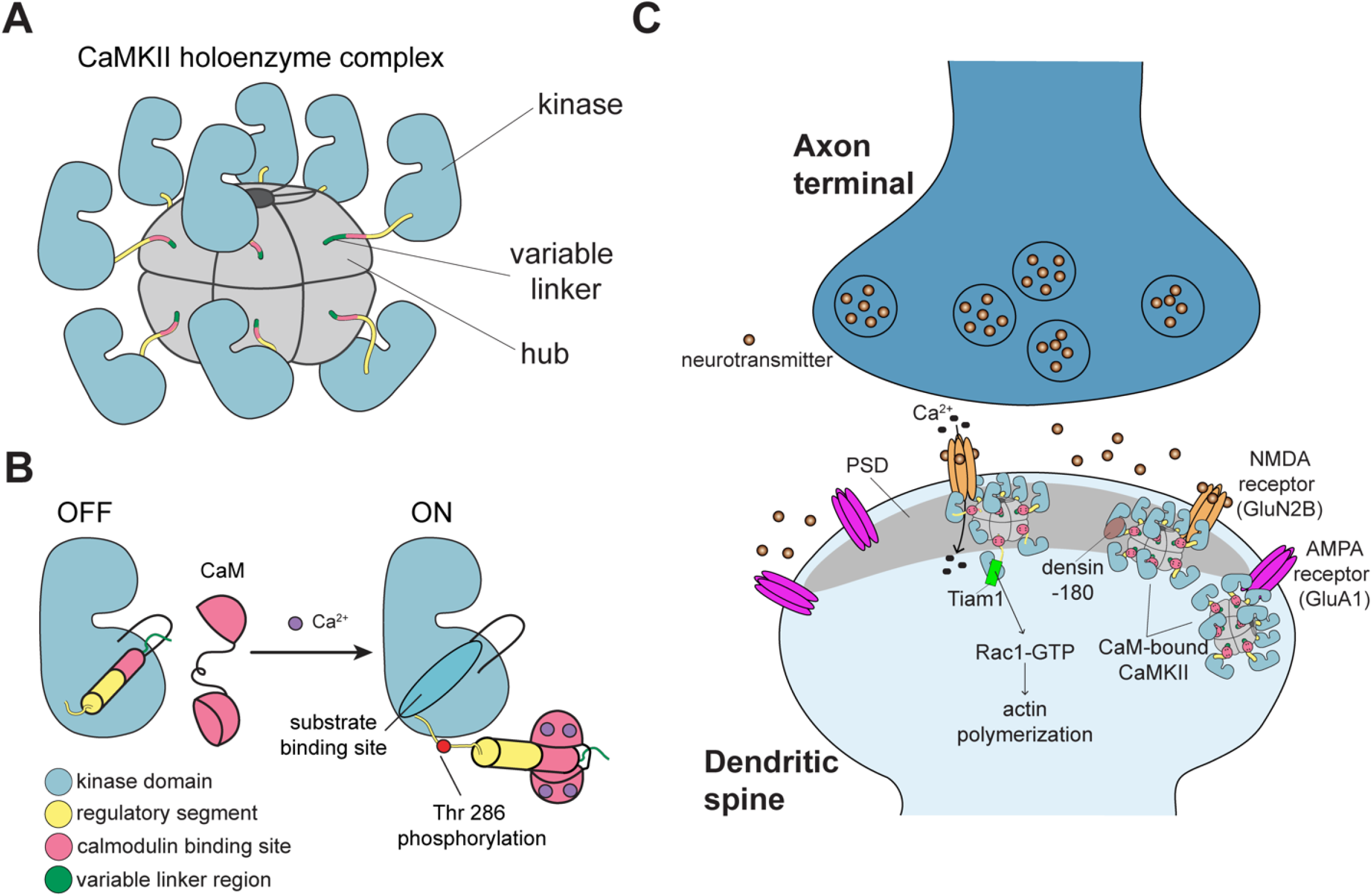
CaMKII architecture and the interaction partners at excitatory synapses. (A) The architecture of a dodecameric CaMKII holoenzyme. (B) Ca^2+^/CaM binding activates CaMKII by competitively binding the regulatory segment thereby freeing the substrate binding site. Active CaMKII autophosphorylates at Thr 286. (C) CaMKII interactions at the excitatory postsynaptic structure, mostly in the post synaptic density (PSD), of the dendritic spine.

Ca^2+^/calmodulin (Ca^2+^/CaM) activates CaMKII by binding to the regulatory segment and thereby freeing the substrate binding site. This well-understood mechanism triggers trans-autophosphorylation of T286, which renders CaMKII constitutively active until T286 is dephosphorylated. CaMKII has also been shown to maintain activity in the absence of Ca^2+^/CaM and T286 phosphorylation by another mechanism that invokes binding partners [6-9], for which we lack a clear mechanism. There are three binding partners (NMDA receptor, Tiam1, and *Drosophila* EAG (*d*EAG)) that have been shown to be activators of CaMKII after the Ca^2+^ stimulus dissipates, with or without T286 phosphorylation [7-10]. The GluN2B subunit of the NMDA receptor is a known substrate of CaMKII [11, 12], and to this point, is the best studied CaMKII activator [13]. Ca^2+^/CaM-activated CaMKII has been shown to form a persistent complex with GluN2B, which locks CaMKII in an active conformation, as long as binding persists [7, 12]. Persistent binding of CaMKII to GluN2B and resultant activation has been explained by a hypothetical model [8]. In this model, GluN2B first binds to Ca^2+^/CaM-activated CaMKII close to the active site (termed S-site), presenting Ser 1303 for phosphorylation. Then, GluN2B dissociates from CaMKII and then re-binds at the base of the CaMKII C-lobe (termed T-site), while freeing the S-site to bind and phosphorylate other substrates. This model has been widely accepted in the field, but to date, there is no structural data supporting it.

Another known activator is Tiam1, a Rac guanine-nucleotide exchange factor (RacGEF), and it is phosphorylated multiple sites by CaMKII. The carboxy tail of Tiam1 also forms a stable complex with CaMKII using a pseudosubstrate sequence (alanine at the phosphorylation site) [9]. The third activator is *d*EAG, a potassium channel in *Drosophila* [10]. Both Tiam1 and *d*EAG behave similarly to GluN2B in that binding maintains CaMKII activation in the absence of Ca^2+^/CaM [9, 10]. A recent structure shows that *d*EAG binds over an extensive surface on the CaMKII kinase domain [14]. On the other hand, densin-180 (LRRC7) is a postsynaptic scaffolding protein. It also forms a stable complex with CaMKII through a pseudosubstrate sequence (isoleucine at the phosphorylation site), but unlike GluN2B or Tiam1, it inhibits kinase activity selectively [15, 16]. Finally, the AMPA receptor subunit GluA1 is a CaMKII substrate but not a known activator [17, 18]. GluA1 phosphorylation by CaMKII is important for synaptic plasticity [19-21].

In the current study, we solved co-crystal structures of the CaMKII kinase domain bound to peptides from GluN2B, Tiam1, densin-180, and GluA1. Using these structures as starting points, we compared molecular dynamics (MD) simulations of the kinase domain in complex with GluN2B, Tiam1, and a previously solved structure with CaMKII bound to an inhibitor, CaMKIIN [22]. Combining this structural information with the biophysical and biochemical measurements has allowed us to clarify important interactions that drive binding and propose a working model for maintaining CaMKII activity in the absence of Ca^2+^/CaM.

## RESULTS

The studies outlined below include co-crystallization of the CaMKII kinase domain bound to peptides from the following CaMKII binding partners: GluN2B (residues 1289-1310), GluA1 (residues 818-837), densin-180 (residues 797-818), and Tiam1 (residues 1541-1559). We also include CaMKIIN in these studies, which is a known endogenous inhibitor of CaMKII [23-25]. Unless otherwise specified, all experiments were conducted in the background of an inactivating CaMKII mutation (D135N). This mutation effectively disables kinase activity by removing the catalytic base while preserving the structural integrity of the kinase. Indeed, an overlay of the wild-type (WT) kinase domain with the D135N kinase domain structure shows very few deviations (root-mean-square deviation (RMSD) = 0.309).

In addition to analysis of the crystal structures, we also performed MD simulations of the kinase domain in complex with extended versions of peptides to assess any contributions from the terminal ends that would not be observed in the crystal structures. All simulations were performed in the NPT ensemble at 300K. One production simulation was performed for the kinase:GluN2B and kinase:Tiam1 complexes, producing trajectories of 2.19 μs and 2.05 μs, respectively. Two independent simulations were performed for the kinase:CaMKIIN complex. These were 0.46 μs and 1.84 μs in length, for an aggregate of 2.3 μs. RMSD and root-mean-square fluctuation (RMSF) for the kinase domain and the peptide were calculated over the course of the trajectories (**Fig. S1A-C**).

### CaMKII kinase domain has a single binding site

We first used isothermal titration calorimetry (ITC) to measure binding affinity of peptides to the CaMKII kinase domain (D135N). We showed that four out of five peptides bound to the kinase with high affinity, whereas GluA1 bound with significantly lower affinity (**Fig. 2A-E**). The K_d_ values for single-site binding ranged from 39 nM to >55 µM (GluN2B K_d_ = 107±47 nM, CaMKIIN K_d_ = 39±24 nM, densin-180 K_d_ = 585±114 nM, Tiam1 K_d_ = 1.1±0.1 µM, GluA1 K_d_ >55 µM) (**Table 4**). From these data, it appears that GluA1 is the outlier whereas the other partners bind with relatively high affinity.

**Figure 2.**
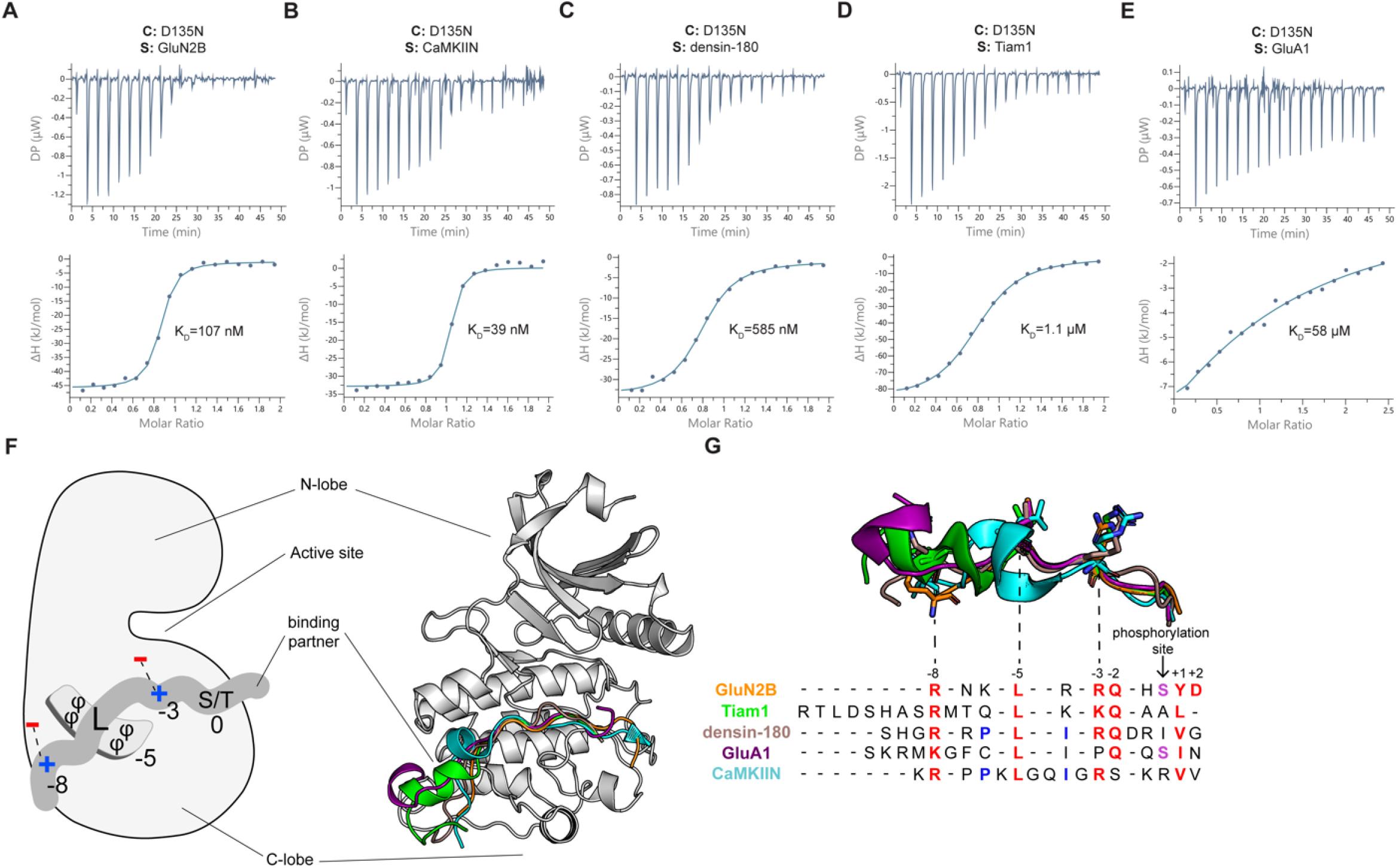
There is a single binding site on the CaMKII kinase domain. (A-E) Representative ITC measurements of D135N kinase domain binding to peptide partners. Contents of the cell (C) and syringe (S) used are listed in the figure panels. The mean K_d_ value from two independent measurements is labeled in the figure. (F) Left: Schematic diagram of the interactions with binding partners. Phi symbol indicates hydrophobic residues. Right: overlay of five cocrystal structures, peptides shown as cartoon in corresponding colors seen in key to the right. (G) The sequence alignment of CaMKII binding partners. Binding partner position numbering is based on the prototypical GluN2B substrate with the phosphorylation site set to zero. Aligned peptide structures are above for reference. Conserved residues are colored red. Phosphorylatable residues at the phosphorylation site are colored purple. Residues involved in a docking event with W214 are colored blue.

We then determined 12 total crystal structures of the CaMKII kinase domain (residues 7-274, lacking the regulatory domain) bound to one of four peptides: GluN2B, GluA1, densin-180, and Tiam1 (**Fig. 2F, S2A-D)**). We resolved 14-19 residues in each peptide. All structures have resolutions within the range of 1.85 – 3.1 Å (**Tables 1-3**). We solved eight structures of the kinase domain bound to the GluN2B peptide (WT and phosphomimetic S1303D) with different ligands in the nucleotide binding pocket (**Tables 1,2**). We also solved structures of the kinase domain bound to peptides of Tiam1, densin-180, and GluA1, with ATP bound or empty nucleotide binding pockets (**Table 3**). Of note, in a structure with both ATP and S1303D GluN2B bound (PDB: 7KL1), there is strong density between the aspartate side chain and gamma phosphate group, indicating the formation of a covalent bond similar to what would be expected for transfer of the gamma phosphate to the carboxylate (**Fig. S2E**).

**Table 1.** GluN2B WT cocrystal structures.

|  | GluN2B/WT<br>kinase/ADP<br>(7UJS) | GluN2B/WT<br>kinase (7UJR) | GluN2B/D135N<br>kinase/ATP<br>(7UJP) | GluN2B/D135N<br>kinase/Hecameg<br>(7UJQ) |
| --- | --- | --- | --- | --- |
| Data collection |  |  |  |  |
| Space group | *P*12<sub>1</sub>1 | *P*12<sub>1</sub>1 | *P*2<sub>1</sub>2<sub>1</sub>2<sub>1</sub> | *P*2<sub>1</sub>2<sub>1</sub>2<sub>1</sub> |
| Cell dimensions |  |  |  |  |
| a, b, c (Å) | 45.01, 65.23, 53.71 | 45.09, 66, 45 | 73.14, 91.42, 91.92 | 73.25, 91.78, 91.77 |
| α, β, γ (°) | 90, 95.26, 90 | 90, 97.61, 90 | 90, 90, 90 | 90, 90, 90 |
| Resolution (Å) | 50 – 2.75 (2.85 – 2.8) | 50 – 1.95 (1.99 – 1.96) | 50 – 2.56 (2.60 – 2.56) | 50 – 2.25 (2.29 – 2.25) |
| *R*<sub>merge</sub> | 0.102 (0.374) | 0.13 (0.485) | 0.232 (0.812) | 0.116 (0.374) |
| Mean *I*/σ*I* | 5 (1.57) | 9.5 (2) | 5 (1.86) | 9.3 (4.36) |
| Completeness (%) | 93.8 (90.9) | 93.8 (82.7) | 100 (99.8) | 96.9 (90.1) |
| Redundancy | 2.6 (2.5) | 2.7 (2.5) | 6.3 (5.4) | 6.3 (5.7) |
| CC<sub>1/2</sub> | 0.985 (0.711) | 0.956 (0.678) | 0.976 (0.572) | 0.99 (0.868) |
| CC* | 0.996 (0.912) | 0.989 (0.899) | 0.994 (0.853) | 0.997 (0.964) |
| Refinement |  |  |  |  |
| Resolution (Å) | 44.86 – 2.75 (2.82 – 2.75) | 33.92 – 1.95 (2.00 – 1.95) | 48-56 – 2.56 (2.62 – 2.56) | 45.93 – 2.25 (2.31 – 2.25) |
| Unique reflections | 7103 (428) | 16929 (1027) | 19465 (1411) | 27648 (1847) |
| *R*<sub>work</sub> / *R*<sub>free</sub> (%) | 22.28/25.56 | 20.93/25.16 | 23.18/28.08 | 21.99/27.11 |
| No. atoms |  |  |  |  |
| Protein | 2253 | 2234 | 4479 | 4507 |
| Water | - | 93 | 45 | 129 |
| Ligand | 27 | 17 | 67 | 52 |
| Ramachadran plot |  |  |  |  |
| In preferred regions (%) | 93.57 | 96.03 | 96.93 | 95.85 |
| In allowed regions (%) | 5.71 | 3.97 | 3.07 | 3.79 |
| Outliers (%) | 0.71 | 0.0 | 0.0 | 0.36 |
| *B*-factors |  |  |  |  |
| Protein | 38.04 | 30.62 | 32.74 | 27.15 |
| Water | - | 33.63 | 18.61 | 21.49 |
| Ligand | 71.96 | 45.15 | 48.55 | 23.01 |
| R.m.s. deviations |  |  |  |  |
| Bond lengths (Å) | 0.0033 | 0.0021 | 0.0037 | 0.0028 |
| Bond angles (°) | 1.2665 | 1.11809 | 1.2648 | 1.2374 |

**Table 2.** GluN2B S1303D cocrystal structures.

|  | GluN2B(S1303D)<br>/D135N<br>kinase/ATP<br>(7KL1) | GluN2B(S1303D)<br>/D135N<br>kinase/ATP<br>(7UJT) | GluN2B(S1303D)<br>/D135N kinase<br>(PDB 7UIS) | GluN2B(S1303D)<br>/D135N<br>kinase/Hecameg<br>(7KL0) |
| --- | --- | --- | --- | --- |
| Data collection |  |  |  |  |
| Space group | *P*2<sub>1</sub>2<sub>1</sub>2<sub>1</sub> | *P*12<sub>1</sub>1 | *P*12<sub>1</sub>1 | *P*2<sub>1</sub>2<sub>1</sub>2<sub>1</sub> |
| Cell dimensions |  |  |  |  |
| a, b, c (Å) | 72.89, 92.13,<br>91.34 | 47.31, 67.25,<br>45.72 | 43.41, 71.43,<br>45.28 | 72.99, 91.29,<br>92.08 |
| α, β, γ (°) | 90, 90, 90 | 90, 94.43, 90 | 90, 97.51, 90 | 90, 90, 90 |
| Resolution (Å) | 50 – 2.4 (2.44 –<br>2.4) | 50 – 2.1 (2.14 –<br>2.1) | 50 – 2.58 (2.64 –<br>2.6) | 50-2.4 (2.44-2.4) |
| *R*<sub>merge</sub> | 0.183 (0.527) | 0.098 (0.297) | 0.162 (0.678) | 0.188 (0.627) |
| Mean *I*/*σ**I* | 5.9 (2.65) | 9.9 (3.51) | 5.8 (1.52) | 4.9 (2.04) |
| Completeness<br>(%) | 99.9 (98.8) | 99.6 (97.3) | 96.9 (92.5) | 99.9 (99.5) |
| Redundancy | 5.7 (5) | 3.3 (2.8) | 3.3 (3.4) | 5.6 (4.9) |
| CC<sub>1/2</sub> | 0.976 (0.740) | 0.993 (0.824) | 0.977 (0.743) | 0.984 (0.652) |
| CC* | 0.994 (0.922) | 0.998 (0.951) | 0.994 (0.658) | 0.996 (0.888) |
| Refinement |  |  |  |  |
| Resolution (Å) | 48.5 – 2.4 (2.46 –<br>2.4) | 38.65 – 2.1 (2.15<br>– 2.1) | 24.37 – 2.58 (2.65<br>– 2.58) | 48.52 – 2.4 (2.46<br>– 2.4) |
| Unique<br>reflections | 23497 (1650) | 15852 (1096) | 7962 (502) | 23396 (1610) |
| *R*<sub>work</sub> / *R*<sub>free</sub> (%) | 20.60/25.75 | 20.34/24.65 | 21.73/ 26.75 | 20.71/25.84 |
| No. atoms |  |  |  |  |
| Protein | 4524 | 2268 | 2207 | 4497 |
| Water | 118 | 96 | 0 | 131 |
| Ligand | 76 | 38 | 0 | 54 |
| Ramachadran plot |  |  |  |  |
| In preferred<br>regions (%) | 97.12 | 95.71 | 95.64 | 96.74 |
| In allowed<br>regions (%) | 2.88 | 3.93 | 4.36 | 3.26 |
| Outliers (%) | 0.0 | 0.36 | 0.0 | 0.0 |
| *B*-factors |  |  |  |  |
| Protein | 28.11 | 28.27 | 43.53 | 25.95 |
| Water | 20.96 | 28.26 | - | 20.24 |
| Ligand | 32.40 | 41.33 | - | 26.08 |
| R.m.s. deviations |  |  |  |  |
| Bond lengths<br>(Å) | 0.0074 | 0.0021 | 0.0030 | 0.0073 |
| Bond angles (°) | 1.4853 | 1.1765 | 1.2691 | 1.4717 |

**Table 3.** Tiam1, Densin-180 and GluA1 cocrystals structures.

|  | Tiam1/D135N<br>kinase/ATP<br>(7UIR) | Tiam1/D135N<br>kinase (7UIQ) | Densin-180/D135N<br>kinase (PDB<br>6X5G) | GluA1/D135N<br>kinase (6X5Q) |
| --- | --- | --- | --- | --- |
| **Data collection** |  |  |  |  |
| Space group | *P*2<sub>1</sub>2<sub>1</sub>2<sub>1</sub> | *P*2<sub>1</sub>2<sub>1</sub>2<sub>1</sub> | *P*12<sub>1</sub>1 | *P*2<sub>1</sub>2<sub>1</sub>2<sub>1</sub> |
| Cell dimensions |  |  |  |  |
| a, b, c (Å) | 43.49, 138.84,<br>154.97 | 43.43, 137.42,<br>156.47 | 36.29, 66.17,<br>61.71 | 42.63, 57.46,<br>107.71 |
| α, β, γ (°) | 90, 90, 90 | 90, 90, 90 | 90, 99.78, 90 | 90, 90, 90 |
| Resolution (Å) | 50 – 3.1 (3.18 –<br>3.13) | 50 – 3.11 (3.17 –<br>3.12) | 50 - 1.85 (1.88-1.85) | 50 – 2.1 (2.18 -<br>2.14) |
| *R*<sub>merge</sub> | 0.287 (0.87) | 0.266 (0.67) | 0.065 (0.325) | 0.179 (0.377) |
| Mean *I*/*σI* | 3.9 (1.47) | 3.9 (1.53) | 9.6 (3.14) | 3.2 (2.21) |
| Completeness<br>(%) | 98 (96.5) | 93.4 (91.9) | 93.6 (86.5) | 96.9 (92.7) |
| Redundancy | 5.6 (4.6) | 4.6 (3.5) | 3.4 (3.3) | 4.9 (4.4) |
| CC<sub>1/2</sub> | 0.975 (0.618) | 0.992 (0.565) | 0.99 (0.829) | 0.992 (0.82) |
| CC* | 0.994 (0.831) | 0.969 (0.831) | 0.997 (0.952) | 0.998 (0.949) |
| **Refinement** |  |  |  |  |
| Resolution (Å) | 33.31 – 3.1<br>(3.183 – 3.103) | 31.54 – 3.11<br>(3.20 – 3.12) | 33.11 - 1.85<br>(1.895-1.85) | 39.32 – 2.14<br>(2.159 – 2.104) |
| Unique<br>reflections | 16512 (1130) | 15535 (1042) | 22043 (1457) | 14475 (798) |
| *R*<sub>work</sub> / *R*<sub>free</sub> (%) | 23.21/27.44 | 25.85/30.92 | 18.71/22.65 | 18.93/23.5 |
| No. atoms |  |  |  |  |
| Protein | 4438 | 4430 | 2259 | 2241 |
| Water | 0 | 0 | 107 | 96 |
| Ligand | 79 | 0 | 23 | 18 |
| Ramachadran plot |  |  |  |  |
| In preferred<br>regions (%) | 93.48 | 91.11 | 97.48 | 95.71 |
| In allowed<br>regions (%) | 5.62 | 8.17 | 2.52 | 4.29 |
| Outliers (%) | 0.91 | 0.73 | 0.0 |  |
| *B*-factors |  |  |  |  |
| Protein | 45.73 | 33.76 | 24.09 | 17.35 |
| Water | - | - | 25.19 | 14.48 |
| Ligand | 62.04 | - | 53.79 | 30.61 |
| R.m.s. deviations |  |  |  |  |
| Bond lengths (Å) | 0.0052 | 0.0068 | 0.0096 | 0.0080 |
| Bond angles (°) | 1.4030 | 1.5793 | 1.6299 | 1.5184 |

**Table 4:**
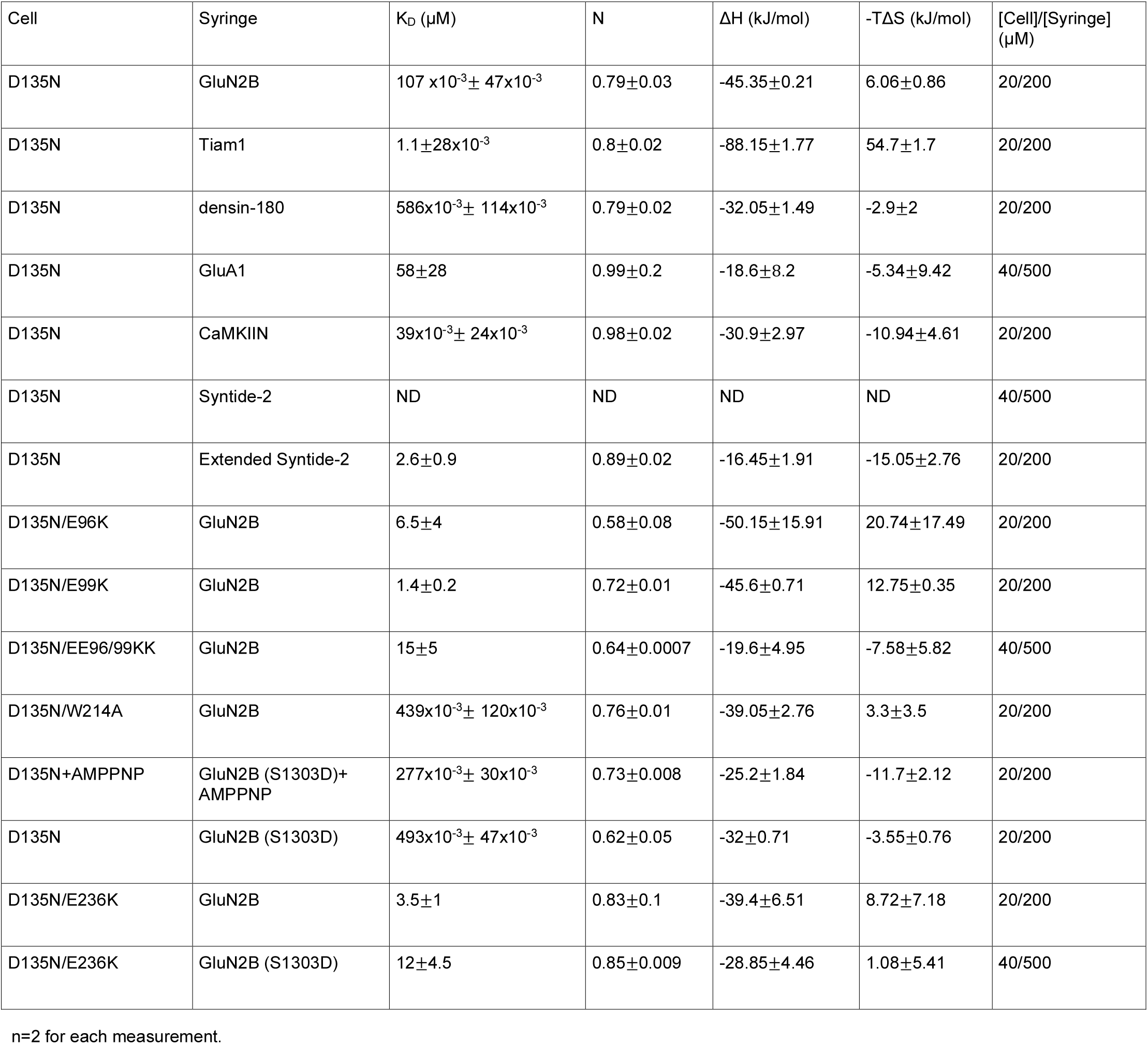
ITC data 1.

Across all structures, we observed that peptides interacted with the same surface on the kinase domain. The overall fold of the kinase domain is similar in all structures. So far, predicting CaMKII interaction partners has been difficult due to conformational heterogeneity. Our structures revealed that several peptides have helical turns, which shifted the register of conserved interactions. In the CaMKIIN structure, there is an internal short helical turn [22]. Tiam1 and GluA1 peptides also adopted short helical turns, however they were shifted to the proximal (N-terminal) end compared to CaMKIIN. GluN2B and densin-180 peptides bound in completely extended conformations with no helical turns. We now provide an updated sequence alignment based on our structural observations (numbering is based on the prototypical GluN2B substrate with the phosphorylation site set to zero and the inserted residues are not counted) (**Fig. 2G**). The two peptides that are substrates (GluN2B and GluA1) have the phosphosite facing the nucleotide binding pocket, such that both peptides are docked at the active site, ready to be phosphorylated. The critical residues of the binding partner mediating this interaction are conserved at positions +1, -2, -3, -5, and -8, which are discussed in detail below (**Fig. 3**).

**Figure 3.**
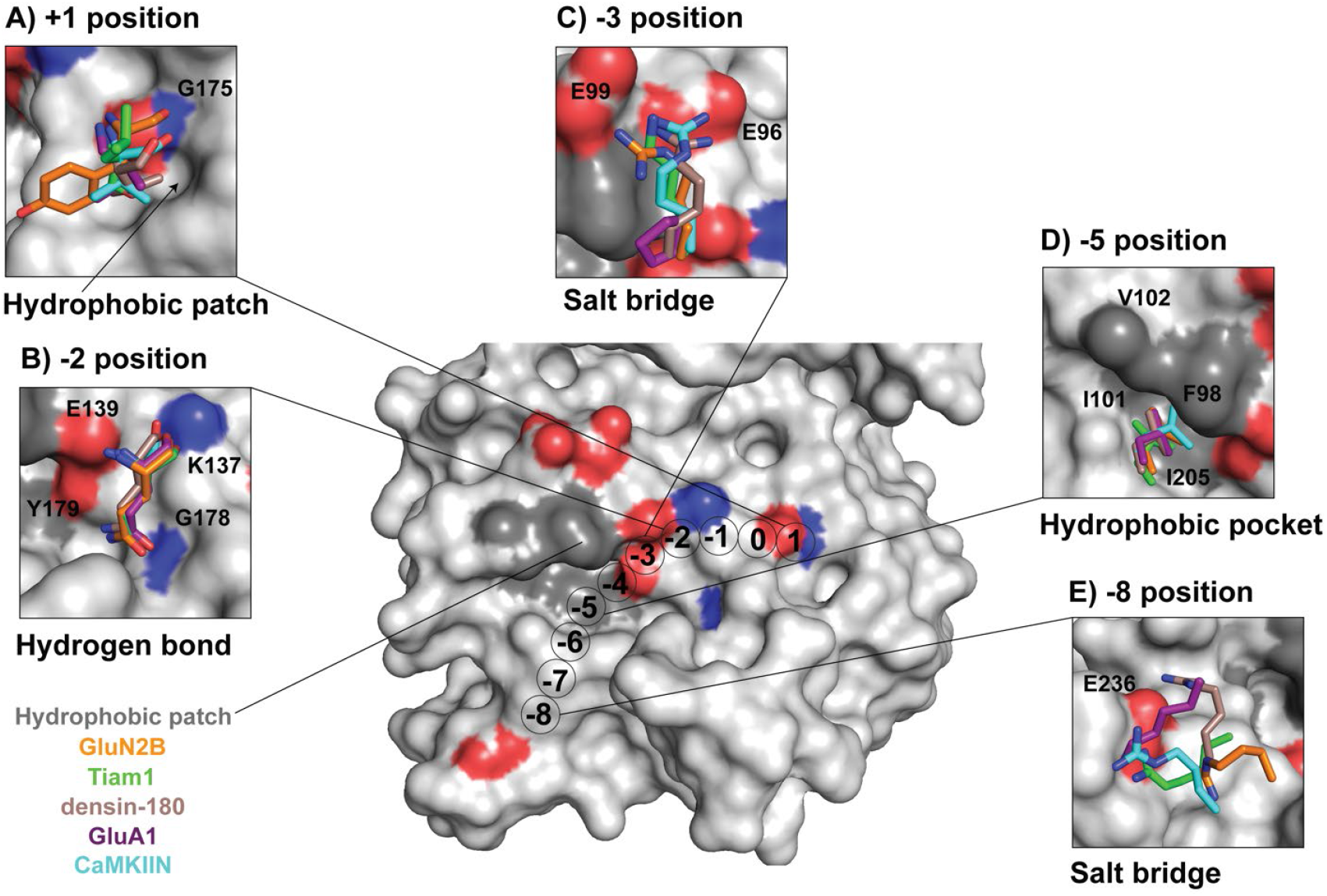
Conserved binding motifs on the catalytic domain surface. Surface representation of the CaMKII kinase domain highlighting residues that mediate interactions with binding partners. Color code matches previous figures (GluN2B: orange, Tiam1: green, densin-180: brown, GluA1: purple, CaMKIIN: cyan). (A) At the +1 position, a hydrophobic patch (arrow) is formed by F173, P177, L185, and Y222. GluA1, CaMKIIN, and densin-180 all have Val or Ile at the +1 position, which are buried in this hydrophobic groove. Backbone atoms of the +1 residue hydrogen bond with the backbone of G175 (highlighted red and blue on the structure). (B) At the -2 position, all interactors have a glutamine except for CaMKIIN, which has a serine. The glutamine sidechain amide oxygen forms a hydrogen bond with the backbone of G178, and the amino group interacts with the sidechain of Y179. The backbone carbonyl interacts with sidechain of K137, and the backbone amino group interacts with E139. (C) At the -3 position, lysine or arginine interact with E96 and E99. The basic residues of interaction partners are positioned 2.4 – 4.2 Å between E96 and E99. GluA1 has a proline at the -3 position, which is flipped away from E96/99. (D) At the -5 position, a conserved leucine across all interactors nestles into a hydrophobic pocket formed by F98, I101, V102, and I205. (E) Lysine or arginine at the -8 position forms a salt bridge with E236.

### Interactions driving binding to the CaMKII kinase domain

#### Hydrophobic residue the +1 position

Our alignment is consistent with previous studies showing that a small hydrophobic residue is preferred at the +1 position [14, 26-29] (**Fig. 2G**). Our structures revealed a hydrophobic patch adjacent to G175 in the kinase domain forms a docking site for small and medium sized hydrophobic residues (valine, isoleucine, and leucine) (**Fig. 3A**). In GluN2B, tyrosine is larger and polar, so the sidechain is not buried in the hydrophobic pocket unlike the other binding partners. In all structures, backbone atoms of the +1 position forms hydrogen bond with the backbone of G175.

In the Tiam1 crystal structure, the +1 leucine does not interact with this groove. We hypothesized that this is because it is the C-terminal residue on the peptide used for crystallization. To test this, we extended the Tiam1 sequence and performed MD simulations, which showed that the +1 leucine does interact with the hydrophobic patch. In the simulations, we observed that the +1 leucine and also the +4 isoleucine interact strongly over the course of the trajectory with an average distance of 3.2 Å (**Fig. S3A**). We compared this to simulations of GluN2B and CaMKIIN, which also show a relatively strong interaction between the +1 residue and the hydrophobic patch (**Fig. S3B, C**). In both GluN2B and Tiam1, the +1 residue fluctuates in and out of the pocket, as shown by the larger distribution of distance from the pocket. Additionally, the +4 residue of GluN2B is also strongly interacting with the pocket similar to Tiam1 whereas the CaMKIIN +2 residue is interacting less.

#### Ionic interactions at the -2 position

Glutamine is commonly found at the -2 position of CaMKII interactors [14, 26-28]. Indeed, all four peptides in our structures have glutamine at -2, allowing us to resolve the important interactions mediated by both the backbone and the sidechain at this position. There is extensive hydrogen bonding between the -2 glutamine and G178, Y179, K137, and E139 (**Fig. 3B**). CaMKIIN has a serine at the -2 position and the side chain is flipped relative to glutamine, which enables tighter interactions with E139 and K137 side chains [22]. This observation is consistent with a previous study, which showed that CaMKIIN binding to CaMKII was reduced when this serine was mutated to alanine [30].

#### Conserved salt bridge at the -3 position

Like many kinases, CaMKII prefers a basic residue at the -3 position [31]. In the crystal structures, all interactors except for the low-affinity binder GluA1 have a basic residue at the -3 position (arginine or lysine), which forms salt bridges with two glutamic acid residues (E96, E99) located on the αD helix of the kinase domain (**Fig. 3C, 4A**) [14, 22, 32, 33]. These salt bridges are not formed with GluA1, because there is a proline at the -3 position. In the GluA1 structure, the sidechain of E96 is flipped away from the peptide compared to the other four structures where it is oriented toward the binding partner (**Fig. 4A**, inset).

**Figure 4.**
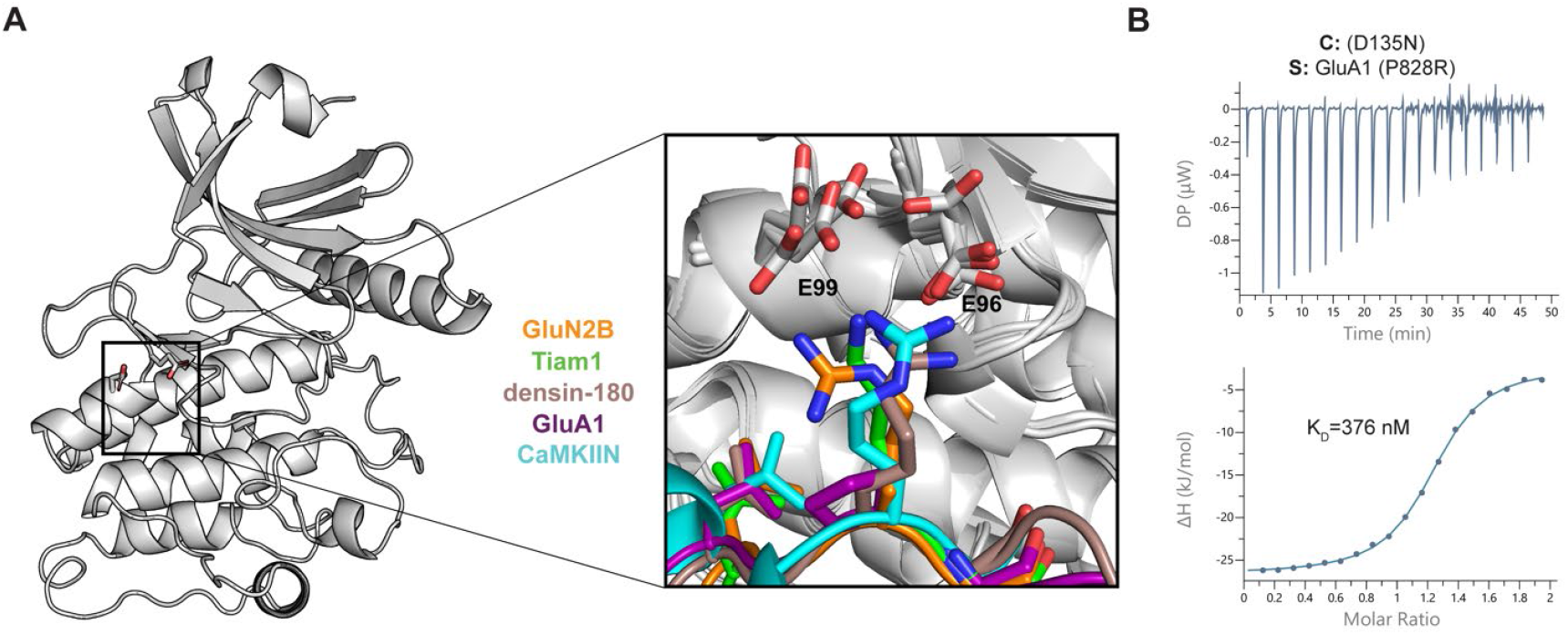
Electrostatic interactions with a basic residue at -3 position facilitate high affinity binding. (A) CaMKII kinase domain shown as a cartoon, E96 and E99 residues are shown as sticks. A zoomed in view of all five co-crystal structures are overlaid to highlight the basic residue at the -3 position (except for GluA1 in purple) interacting with the two glutamic acids in the kinase domain. (B) ITC data between D135N CaMKII kinase domain (cell) and GluA1 with P828R mutation (syringe). The mean K_d_ value is from two independent measurements.

We hypothesized that the reason for GluA1’s lower affinity was due to the lack of these salt bridges. To test this, we measured binding of a mutant GluA1 peptide where the -3 proline was changed to an arginine (GluA1 P828R). This mutation increased binding >140-fold to K_d_ = 376 nM (**Fig. 4B**). To further validate this, we created single and double charge reversal mutations (E → K) and compared the behavior of GluA1 P828R to a high affinity interactor, GluN2B. Single mutants (E96K or E99K) decreased the affinity to both GluA1 P828R and GluN2B by 7-to 65-fold, whereas the double mutant (E96/99K) decreased affinity by 75-to 140-fold (**Fig. S4, S5, Tables 4, 5**). These data indicate the importance of these salt bridges in determining binding affinity.

**Table 5:** ITC data 2.

| Cell | Syringe | K<sub>D</sub> (μM) | N | ΔH (kJ/mol) | -TΔS (kJ/mol) | [Cell]/[Syringe] (μM) |
| --- | --- | --- | --- | --- | --- | --- |
| D135N+AMPPNP | CaMKIIN+ AMPPNP | 22x10<sup>-3</sup>±4x10<sup>-3</sup> | 0.91±0.007 | -40.7±0.14 | -2.35±0.52 | 8/80 |
| D135N/E96K +AMPPNP | CaMKIIN+ AMPPNP | 397x10<sup>-3</sup>± 15x10<sup>-3</sup> | 0.79±0.008 | -43±2.12 | 7.06±2.19 | 20/200 |
| D135N/E99K +AMPPNP | CaMKIIN+ AMPPNP | 87x10<sup>-3</sup>±12x10<sup>-3</sup> | 0.86±0.004 | -58.8 | 19.15±0.35 | 20/200 |
| D135N/EE96/99KK +AMPPNP | CaMKIIN+ AMPPNP | 1.4±0.3 | 0.58±0.01 | -103.8±7.43 | 70.65±7.7 | 20/200 |
| D135N/E236K | CaMKIIN | 825x10<sup>-3</sup>±6x10<sup>-3</sup> | 0.87±0.02 | -29.65±0.78 | -4.53±0.76 | 20/200 |
| D135N/W214A | CaMKIIN | 2±0.2 | 0.97±0.007 | -25.05±0.21 | 6.57±0.4 | 40/500 |
| D135N/W214A | Tiam1 | 10±4 | 1.56±0.1 | -43.7±5.8 | 15.55±6.72 | 40/500 |
| D135N/E236K | Tiam1 | ND | ND | ND | ND | 40/500 |
| D135N/W214A | densin-180 | 2±0.7 | 0.98±0.006 | -10.07±2.03 | -21.9±2.83 | 20/200 |
| D135N/E236K | densin-180 | 10±7 | 1.36±0.25 | -11.14±2.07 | -17.25±3.75 | 35/500 |
| D135N | GluA1 (P828R) | 376x10<sup>-3</sup>±110x10<sup>-3</sup> | 1.17±0.04 | -23.4±1.41 | -12.7±2.12 | 20/200 |
| D135N/E96K | GluA1 (P828R) | 2.7±1.8 | 1.06±0.14 | -8.08±2.54 | -23.55±4.31 | 20/200 |
| D135N/E99K | GluA1 (P828R) | 5±0.5 | 1.33±0.4 | -15.9±0.57 | -13.8±0.71 | 20/200 |
| D135N/EE96/99KK | GluA1 (P828R) | 28±3 | 1.22±0.16 | -4.52±0.79 | -21.05±1.06 | 40/500 |

n=2 for each measurement.

We also wanted to investigate the role of E96/99 in ATP binding. E96 is well-conserved across kinases and known to be important for ATP binding [33]. Almost 50% of kinases have either an aspartate or glutamate at this position (E96), which might explain the basophilic kinase substrate preference [34]. In structures where ATP/Mg^2+^ is bound (GluN2B, Tiam1), the side chain of E96 interacts with the hydroxyl groups of ribose in ATP (**Fig. S6A**). In MD simulations with CaMKIIN, interactions with ATP were very different. We observed a unique interaction at the -3 position, which resulted in a conformation that distorted ATP binding (**Fig. S6B**). MD trajectories predict that CaMKIIN binding is facilitated by E96 in the presence of ATP, whereas E99 plays essentially no role. We performed ITC measurements comparing binding of CaMKIIN to the kinase domain with charge reversal mutations (E96K, E99K) in the presence of an unhydrolyzable ATP analogue (AMP-PNP). Consistent with the prediction from the simulation, the affinity of E99K was only 2-fold weakened while E96K weakened by 10-fold (**Fig. S4, S6C-F, Table 5**). Further efforts will need to interrogate the role of ATP coordination in CaMKIIN inhibition. E99 is not as well-conserved, even across other CaM-kinases (**Fig. S7A-C**), indicating that it may be more important for substrate specificity and potentially compensate if E96 is mutated [35].

#### Hydrophobic interaction at the -5 position

All CaMKII interactors studied here have a conserved leucine at the -5 position, which fits into a hydrophobic pocket on the kinase domain comprised of F98, I101, V102, and I205 (**Fig. 3D, 5A**). In these structures, leucine is 3.3-4.5 Å from the four hydrophobic residues, indicating a tight interaction as compared by measuring the distance between the closest carbon atoms in each residue. In the CaMKIIN structure, a turn motif, facilitated by two glycines, orients leucine at the -8 position into this pocket to make it nominally at the -5 position [22]. In simulations of GluN2B, Tiam1, and CaMKIIN, this interaction also demonstrates high structural integrity, with pairwise RMSD below 2 Å (**Fig. 5B**).

**Figure 5.**
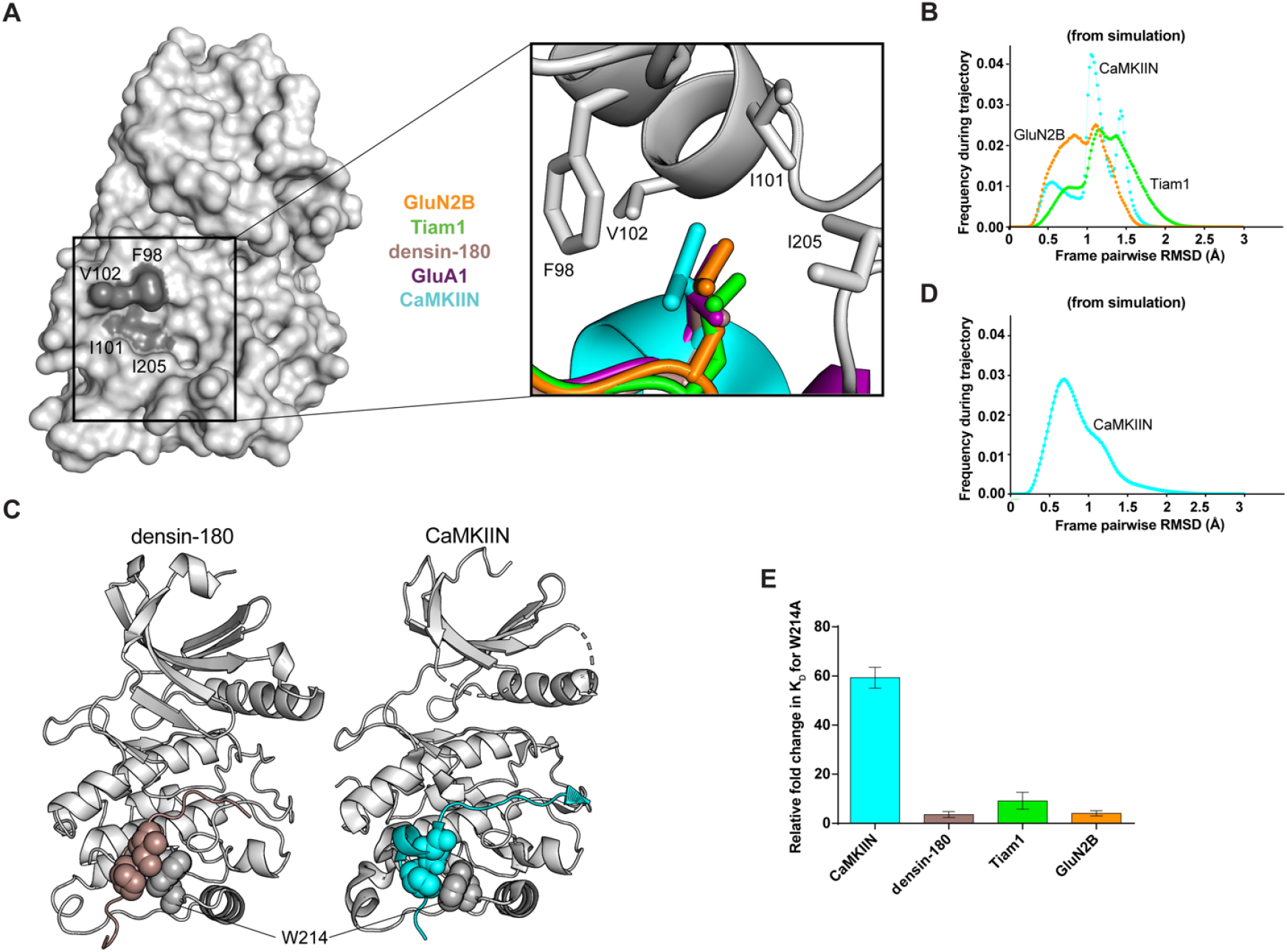
Hydrophobic interactions mediate binding. (A) Surface representation of CaMKII kinase domain with residues forming the hydrophobic pocket labeled (dark gray). Inset: overlay of leucine residues from all co-crystal structures bound in the hydrophobic pocket. (B) Histograms of RMSD from MD simulations between every pair of trajectory frames for F98, I101, V102, and I205 with -5 peptide leucine. (C) Crystal structures with sphere representation of the isoleucine and proline or leucine residue of densin-180 (brown) and CaMKIIN (cyan) docking onto W214 (gray) of kinase domain. (D) RMSD histogram from MD simulations for W214 interacting with isoleucine and proline of CaMKIIN. (E) K_d_ values were extracted from ITC data and relative fold changes were calculated by dividing the observed K_d_ from the mutant (W214A) by the D135N kinase domain. Standard deviations from the mean are shown as error bars for duplicate measurements.

#### Docking site mediated by W214

In both densin-180 and CaMKIIN structures, the sidechains of proline and isoleucine (highlighted blue in **Fig. 2G**) pack against W214 on the kinase domain (**Fig. 5C**). For densin-180, the guanidino group of R808 at the -7 position hydrogen bonds with the backbone carbonyl group of W214 and the sidechain of Q224. In simulations of CaMKIIN, the pairwise RMSD calculated for W214 interacting with isoleucine and proline below 1.5 Å, indicating a strong and persistent interaction over the course of the trajectory (**Fig. 5D)**.

We measured binding to a kinase domain mutated at this tryptophan (W214A) to discern its contribution to these interactions. The W214A mutation significantly disrupted CaMKIIN binding by lowering the affinity 60-fold, whereas the effects on the other binding partners were not as severe (4-to 10-fold decreased affinity) (**Fig. 5E, Fig. S8A-D, Tables 4, 5**). CaMKIIN docking onto W214 likely stabilizes its helical motif, which properly orients the -5 leucine into the hydrophobic pocket. When W214 is mutated, the -5 leucine interaction is unlikely to be maintained in CaMKIIN, which results in disrupted binding. Densin-180 does not have a helical motif, which may explain why the W214A mutant does not have a drastic effect on binding. CaMKIIN has been characterized as a CaMKII specific inhibitor despite the fact that CaMKII has high similarity to the CaMK-family kinases. This specificity is likely driven by the presence of this tryptophan residue as the other family members have tyrosine or methionine at this position (**Fig. S7A**).

#### Salt bridge at the -8 position

We observed a novel electrostatic interaction between a conserved basic residue at the -8 position (arginine or lysine) and a glutamate (E236) at the base of the C-lobe (**Fig. 3E, 6A**). This has not been previously described likely because this position is quite far from the phosphorylation site (∼30 Å), and substrate alignments have not been accurate enough to highlight the conservation at this position. We tested the effect of mutating E236 to lysine (E236K) on binding affinity. The E236K mutation completely abolished Tiam1 binding (**Fig. 6B, Fig. S8E, Table 5**). For densin-180, CaMKIIN, and GluN2B, E236K binding affinity was reduced ∼17-fold (K_d_ = 10±7 µM), ∼21-fold (K_d_ = 825±6 nM) and ∼35-fold (K_d_ = 3.5±1 µM), respectively (**Fig. 6B, Fig. S8F-H, Tables 4, 5)**. Previous work noted that E236K did not significantly affect GluN2B binding in biochemical assays [8], however, in that measurement low micromolar binding would not be discernable from high nanomolar binding as in our ITC measurements.

**Figure 6.**
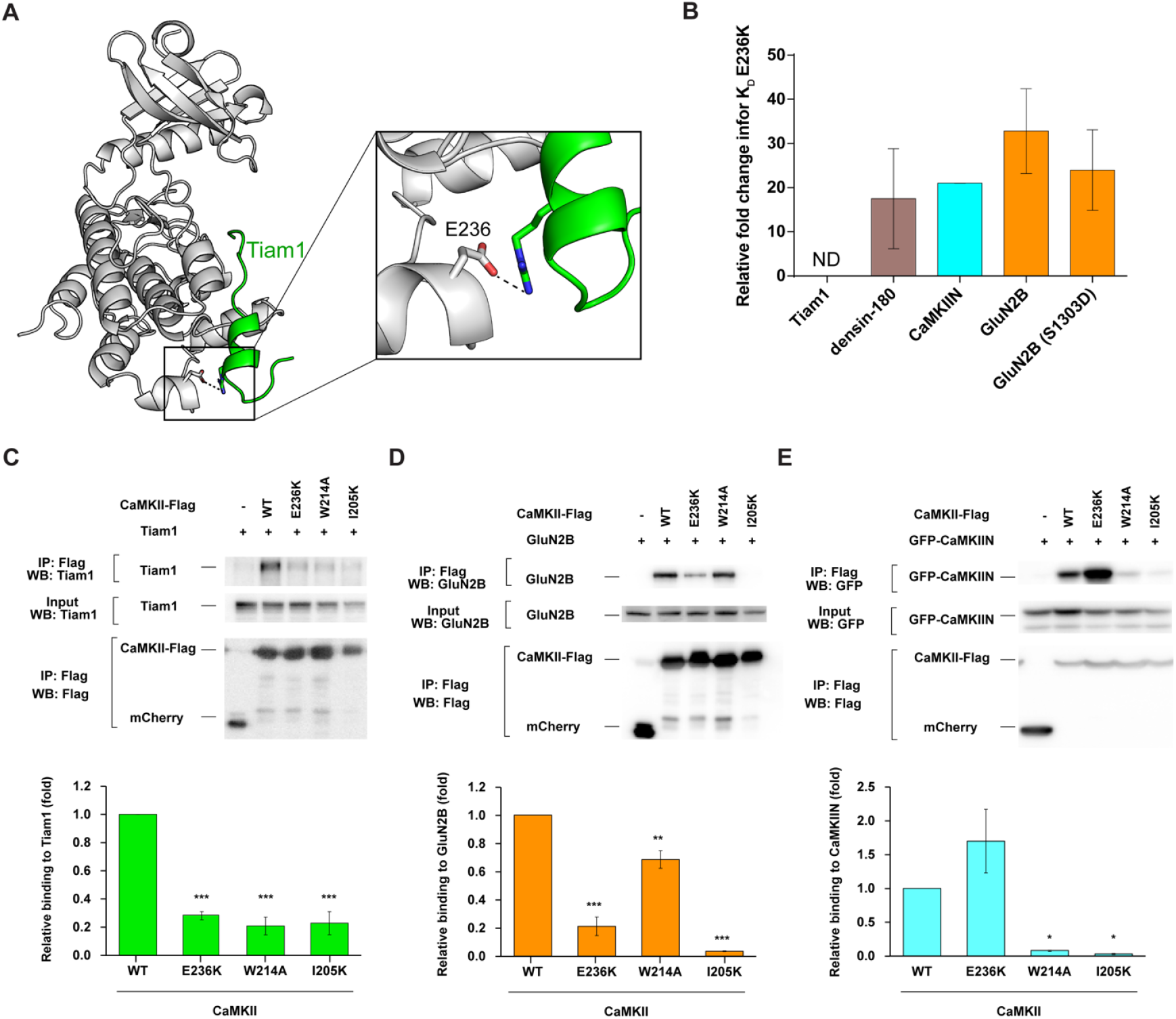
Electrostatic interaction with E236 facilitates binding. (A) View of the interaction between CaMKII E236 (gray) and Tiam1 R1549 (green). (B) K_d_ values were extracted from ITC data and relative fold changes were calculated by dividing the observed K_d_ from the mutant (E236K) by the D135N kinase domain. Standard deviations from the mean are shown as error bars for duplicate measurements. (C-E) Effects of CaMKII mutations (E236K, W214A, and I205K) on interactions with Tiam1, GluN2B, and CaMKIIN. HEK293T cells were co-transfected with Flag-tagged CaMKII variants and Tiam1-mGFP, GluN2B or EGFP-CaMKIIN. Cell lysates were immunoprecipitated with Flag antibody, and samples were immunoblotted with Tiam1, GluN2B, GFP, and Flag antibodies. Representative blots are shown in upper panels. Quantification of the co-immunoprecipitation from 3 or 4 independent experiments were shown in a graph of (C) Tiam1 (n=3) (D) GluN2B (n=3) (E) CaMKIIN (n=4). Error bars indicate standard error of the means. The amount of co-immunoprecipitated Tiam1, GluN2B or CaMKIIN was normalized by the amount in cell lysate and immunoprecipitated CaMKII. *p < 0.05, **, < 0.01 and ***, < 0.001, compared to WT CaMKII; one-way ANOVA with the Shaffer’s post hoc test comparisons. WT, wild type; IP, immunoprecipitation.

We wondered whether substrate phosphorylation would alter this interaction that is so far from the active site. To test this, we measured the affinity between the E236K kinase domain and the phosphomimetic GluN2B (S1303D, S→D at position 0). There was very little effect on the S1303D GluN2B binding to D135N (K_d_ = 493±47 nM) (**Fig. S9A**), whereas affinity for the E236K mutant was reduced ∼24-fold (K_d_ = 12±4.5 µM) (**Fig. 6B, Fig. S9B, Table 4**). This indicates that E236K significantly weakens binding to S1303D GluN2B.

### Generalizing CaMKII binding partners

We mined the curated database of CaMKII phosphorylation sites (n=418) for consistencies with our updated alignment and found additional similarities (**Fig. S7D**, www.phosphosite.org) [40]. Hydrophobic residues (I, V, L, M, F) are found at +1, Q is found at -2, and R is overwhelmingly found at -3, but also at -2 and -4. Smaller hydrophobic residues (L, V, I, M) are found at -5 and - 6. Generally, the minus direction disfavors P, D, and E. In contrast, acidic residues D and E are highly favored at +2. GluN2B has D at the +2 position, and we note an interaction with K56 in several structures, which was also observed in the *d*EAG-bound structure (PDB: 5H9B) [14]. In addition, S, which may serve as a second phosphorylation site, is strongly disfavored at nearly every position. The Phosphosite Plus database only allows analysis out to +7/-7 positions so we could not compare residues at the -8 position and beyond. The actual -8 position may be farther than anticipated from alignments due to secondary structural elements. Overall, these features are consistent with important residues identified in our structures.

### Interrogating CaMKII interactions with full-length binding partners

In order to assess the importance of the residues on CaMKII identified in the crystallographic analysis, we used a pull-down assay to assess the effect of CaMKII mutations on interactions with full-length binding partners. We co-transfected HEK293 cells with FLAG-tagged full-length CaMKIIα and either full-length Tiam1, GluN2B, or GFP-CaMKIIN. Binding was assessed by western blot following immunoprecipitation with an anti-FLAG antibody. Immunoprecipitation showed a remarkable loss of binding for both Tiam1 and GluN2B when E236 is mutated, similar to our ITC data (**Fig. 6C, D**). E236K CaMKII resulted in better binding to CaMKIIN compared to WT CaMKII in HEK293 cells (**Fig. 6E**). Our ITC results showed tight binding between the E236K CaMKII kinase domain and CaMKIIN (825 nM), and it is known that the E236K mutation disrupts regulatory segment binding [32], thereby facilitating more CaMKIIN binding to E236K compared to WT since the binding site is more available. The far salt bridge interaction mediated by E236 facilitates high affinity binding to CaMKII and may also be critical for CaMKII specificity as it is not conserved across other CaMK-family kinases.

The immunoprecipitation experiments also corroborate our peptide binding data that show the importance of hydrophobic interactions. Indeed, mutating the hydrophobic pocket at the -5 position (I205K) showed significant loss of CaMKII binding to Tiam1, GluN2B, and CaMKIIN (**Fig. 6C-E**). The W214A mutation also disrupted binding to Tiam1, GluN2B, and CaMKIIN. The pull-down assay revealed a significant effect on Tiam1 binding to W214A, which is in agreement with ∼9-fold lower affinity from ITC measurements (**Fig. 6C**). The W214A mutation slightly weakened the interaction with full-length GluN2B, and we observed 4-fold lower affinity in our ITC measurement using GluN2B peptide (**Fig 6D**). Consistent with our ITC data, CaMKIIN binding to W214A was remarkably weakened compared to WT (**Fig. 6E**). CaMKIIN is a high affinity inhibitor of CaMKII, and its specificity likely arises from this W214 docking site which is not present in other CaMK-family members (**Fig. S7A, C**). In this way, W214 might serve as a docking site for CaMKII-specific interactions. Additionally, a previous study suggested that the pantothenate segment of Coenzyme A packs against W214 of CaMKII to mediate this interaction [41].

### The αD helix rotates outward to accommodate binding partners

CaMKII is unique among protein kinases in that it does not require phosphorylation in the activation loop. In fact, CaMKII structures have highlighted that the autoinhibited conformation of CaMKII actually resembles many other kinases’ active conformations [36, 37]. Upon activation, the αD helix in CaMKII undergoes a 45° rotation outward [33, 36, 38]. In the autoinhibited conformation, the regulatory segment stabilizes the αD helix in the inward conformation [36].

We then compared the structures of autoinhibited kinase (PDB:2VZ6, residues 13-300) [33], the uninhibited kinase (PDB:6VZK) [39], and the kinase bound to GluN2B. It is clear from this comparison that that the αD helix rotates outward to accommodate GluN2B binding, just as it does when the regulatory segment is removed (**Fig. 7A**). This is also true for Tiam1, GluA1, densin-180, and CaMKIIN [22]. The rotation of the αD helix effectively presents the hydrophobic patch in a competent position for interacting with the binding partner. The αD helix houses three of the four residues that make up the hydrophobic pocket for the -5 leucine to dock in, which is likely a large contribution to an observed gain in kinase domain stability upon substrate binding [6] (**Fig. S10A**), similar to what we have previously observed with regulatory segment binding, which is mediated by L290 in the regulatory segment [39].

**Figure 7.**
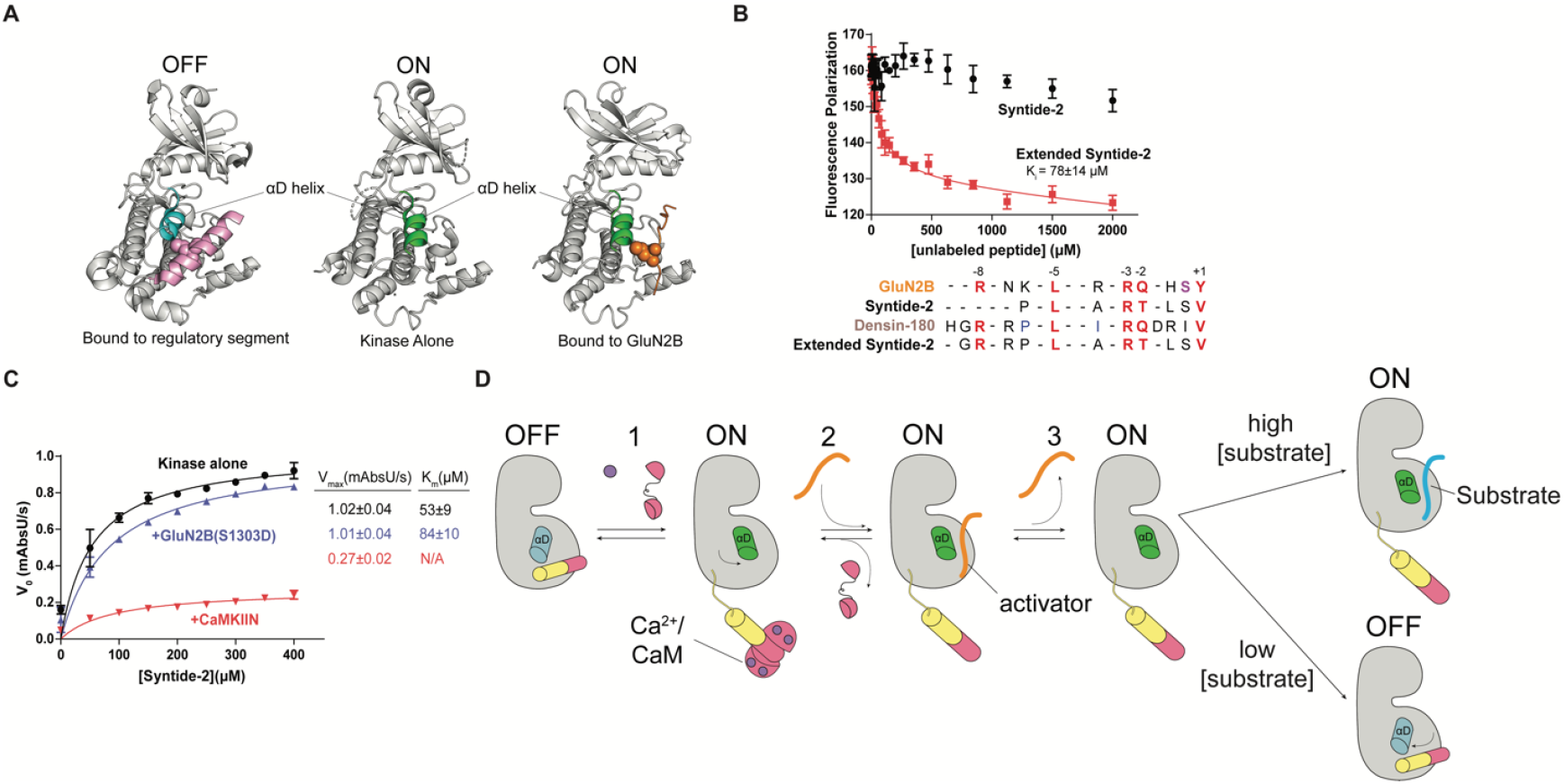
GluN2B acts as a competitive inhibitor on CaMKII. (A) Comparison of the αD helix between crystal structures of the autoinhibited with regulatory segment (pink) bound (left, PDB:2VZ6), uninhibited kinase with no regulatory segment (middle, PDB:6VZK), and active kinase with GluN2B (orange) bound (right, PDB:7UIS). The αD helix goes from rotated inward in the off-state (blue) to rotated outward 45° in the on-state (green). (B) Competition assay against GluN2B using Syntide-2 (black) and extended version of Syntide-2 (red). Sequence alignment of GluN2B, Syntide-2, densin-180, and the extended Syntide-2 is shown below the graph. For clarity, the alignments start at the +1 position (C-terminal end) and contain the -8 basic residue (N-terminal end). Full sequences used are listed in the Methods section. (C) Coupled kinase assay results with kinase alone, in the presence of GluN2B and CaMKIIN. The significance of the change in K_m_ values is significant using unpaired t test with Welch’s correction, p=0.0208. (D) Proposed model of maintaining CaMKII activity by binding to a high affinity activator. CaMKII binds Ca^2+^/CaM and is activated, and the αD helix rotates out (Rxn 1). A high affinity activator binds to the substrate binding site (Rxn 2). When the Ca^2+^ signal dissipates, CaM dissociates from CaMKII, but the activator remains bound, competing with the regulatory segment. The activator dissociates, but the αD helix remains in the active conformation (Rxn 3). In high [substrate], substrate will bind and be phosphorylated. In low [substrate], either the activator will rebind, or the αD helix will rotate back in to accommodate regulatory segment binding.

### Testing the S- and T-site binding model

Since our structures revealed that all interactors bound to the same single site on the kinase domain, we wanted to directly test the idea of the 2-site (S- and T-site) binding model [8]. Previous studies have shown that GluN2B peptide could not be competed off by Syntide-2 even at low millimolar concentrations, which we also observed (**Fig. 7B**) [12]. As noted by Colbran and colleagues [12], a sequence alignment of Syntide-2 compared to other substrates reveals that the Syntide-2 peptide is shorter at the N-terminus and is therefore missing the -8 position basic residue necessary for mediating the far salt bridge. To test whether this is the reason Syntide-2 could not compete GluN2B, we created an extended version of Syntide-2 by adding three residues to the N-terminus, mimicking densin-180 (**Fig. 7B**, inset). The extended version of Syntide-2 did compete off GluN2B (K_i_ ∼ 78±14 μM) (**Fig. 7B**). This result shows that Syntide-2 and GluN2B occupy the same binding site, indicating that there are not two separate binding sites on the kinase domain. When we tried to measure affinity for Syntide-2 peptide using ITC, we did not see any appreciable binding despite using protein/peptide concentrations high enough to characterize an interaction with a K_d_ of several hundred micromolar (**Fig. S11**). When we tested the extended version of Syntide-2 in ITC, we obtained a K_d_ of 2.6±0.9 µM **Fig. S11**, providing further evidence that this far salt bridge at the -8 position facilitates tight binding.

A conundrum thus arises: if there are not two distinct binding sites, how is CaMKII activity maintained by specific binding partners like GluN2B? The mode of activation must invoke the phosphorylated form of GluN2B since this is observed under activating conditions. In a canonical kinase reaction, the kinase first binds substrate and ATP, then transphosphorylation occurs, and finally the substrate is released and ADP is replaced by another ATP molecule. We measured the affinity of the phosphomimetic GluN2B peptide (S1303D) (K_d_ = 493±47 nM) and observed only a ∼5-fold reduction in affinity compared to WT GluN2B (K_d_ = 107±46 nM) (**Fig. S9A, 2B**). In the presence of AMPPNP, there is also tight binding of S1303D GluN2B (K_d_ = 277±30) (**Fig. S9C**). We were also able to crystallize the kinase domain bound to GluN2B S1303D, which showed the same conformation as bound to WT GluN2B (backbone RMSD 0.266 Å) (**Fig. S10B**). We hypothesized that phospho-GluN2B would act as a competitive inhibitor for Syntide-2 due to this tight binding. We performed kinase assays using Syntide-2 as a substrate in the presence of saturating GluN2B S1303D. As a control, we also measured activity against Syntide-2 in the presence of the CaMKIIN. Indeed, high levels of CaMKIIN completely inhibit Syntide-2 phosphorylation, whereas GluN2B acts as a competitive inhibitor, exemplified by comparable V_max_ values but an increased K_m_ value (**Fig. 7C**). These data indicate that GluN2B and Syntide-2 bind to the same site.

### A proposed model for persistent CaMKII activity

Based on our data, we propose a mechanistic model to explain the observations of prolonged CaMKII activity in the presence of specific binding partners. Let’s consider the mode of activation by GluN2B as the activator shown in **Figure 7D**. As described above, upon Ca^2+^/CaM binding and release of autoinhibition, the kinase domain undergoes a conformational change where the αD helix rotates out and the kinase is in the ‘on’ state (**Fig. 7A**). Following dissociation of GluN2B, a conformational change in the αD helix is required for the regulatory segment to re-bind but not for a new substrate to bind. For this reason, we hypothesize that under conditions of high substrate concentrations, substrate binding will be faster than regulatory segment re-binding. In conditions where the activator concentration is high, activator rebinding will be favorable since this high affinity binder will have a relatively high on-rate and low off-rate.

## DISCUSSION

CaMKII recognizes a broad range of interaction partners, yet the structural details driving these interactions had not been elucidated. We addressed this by solving co-crystal structures of four binding partners bound to the CaMKII kinase domain, which allowed us to highlight the interactions mediating binding. The minimum consensus sequence for CaMKII targets is R-X-X-S/T [42, 43], however, our results have expanded upon this understanding. If we incorporate our structural observations, the full consensus sequence is: R/K-X-X-ϕ-X-R/K-X-X-S/T-ϕ. We postulate that if all positions are satisfied as shown here, the binding affinity will be high nanomolar to low micromolar. Interaction partners lacking 1-2 of these positions will have poorer affinities but may still be viable interactors, since a typical kinase substrate has an affinity of several hundred micromolar [44]. For example, GluA1 lacks a basic residue at the -3 position which yields an affinity of >55 µM. The biological ramifications of the weak interaction between CaMKII and GluA1 will need to be further investigated – where localization of CaMKII to this receptor by another factor or avidity effects of holoenzyme localization may be driving GluA1 phosphorylation *in vivo* [45]. Our observation might explain the molecular basis of a recent study that reported lower levels of S831 phosphorylation in the hippocampus [46]. We report GluA1 as a low-affinity substrate, which would explain why densin-180 was interpreted as a selective inhibitor since GluA1 would not be strong enough to compete with densin-180 binding [16]. Given the large size and scaffolding role of densin-180 (180 kDa) compared to CaMKIIN, the densin-180 interaction might be more important for CaMKII localization than widespread inhibition.

There has been a lack of mechanistic clarity in understanding how certain CaMKII interactors lead to prolonged activation in the absence of calcium. In 2001, without a crystal structure to guide an understanding, a set of extraordinary data revealed that GluN2B binding yielded autonomous CaMKII activity [7]. This work came on the heels of an extensive mutagenesis study mapping how the regulatory domain interacts with the surface of the kinase [32]. One of the findings from this work was that the mechanism of autoinhibition is distinct from substrate binding, especially when residues far from the active site were tested. To orient thinking about these interactions, residues close to the active site were termed the S-site (for substrate binding) and residues far from the active site were termed the T-site (for Thr286 binding) [38, 47]. In 2006, these reference points were implemented into a 2-site binding model to explain this autonomous activity with GluN2B binding [8].

We were well-poised to systematically test the S- and T-site model using biochemistry and structural biology. In the described model, F98 is part of the S-site and I205 is part of the T-site [8]. Conversely, we find that all interactors, including activators, bind to the CaMKII kinase domain using a continuous single site across the surface of the kinase domain [14, 22]. In these structures, F98 and I205 (along with I101 and V102) make up a single hydrophobic pocket that houses the -5 leucine residue, first described by Kuriyan and colleagues as ‘docking site B’ (**Fig. 3D, 5A**) [22]. So, F98 and I205 are performing the same role, instead of comprising separate sites. A large body of evidence for the S- and T-site model has been based on experiments that compare a short synthetic substrate Syntide-2 (lacks a basic residue at the -8 position) to another substrate AC3 (contains a -8 basic residue). Syntide-2 has been deemed an S-site binder whereas AC3 binds to both the S- and T-site. This distinction stemmed from data showing that if CaMKII is bound to Syntide-2 or AC3 and then GluN2B is added, GluN2B binds the CaMKII-syntide-2 but not the CaMKII-AC3 [8]. These data were interpreted such that GluN2B was able to bind to the T-site while Syntide-2 remained bound at the S-site. Based on our data, an updated interpretation is that GluN2B competes off Syntide-2 because GluN2B binds CaMKII with >1000-fold higher affinity.

CaMKII has high affinity (submicromolar to low micromolar) for activator peptides GluN2B, Tiam1, and *d*EAG [14]. This is somewhat unique, since typically these interactions are in the K_d_ range of 200 µM [44]. We postulate that these high affinity interactors allow the interaction partners to kinetically compete with regulatory segment rebinding by stabilizing the active conformation with the αD helix rotated outward. In conditions where substrate concentration is high, CaMKII will phosphorylate substrate because the kinase is in the active conformation and is in the proper conformation to interact with substrates. The data presented here provide clarity on the interactions between CaMKII and its binding partners, which will be crucial in guiding biological experiments to assess the downstream effects of specific interactions. Future experiments will necessarily invoke the complexities of the CaMKII holoenzyme structure. CaMKII-activator binding provides a mechanism to maintain CaMKII activity in the absence of Ca^2+^. It is intriguing to consider other enzymes that might be modulated in a similar way by high affinity binding partners as a novel mechanism for prolonged activation in the absence of other stimuli.

## MATERIALS AND METHODS

### Cloning

All CaMKIIα variants were expressed with an N-terminal His-SUMO tag in a pET vector. For mammalian expression, CaMKIIα was expressed with a C-terminal Flag tag in pDEST12.2 vector [9]. Tiam1-mGFP was subcloned in a pCAGGS [9], GluN2B was in pGW-CMV (Kim et al., 1996, https://doi.org/10.1016/S0896-6273(00)80284-6), CaMKIIN-GFP was in pEGFP vector [25], and mCherry-Flag was in pDEST12.2 vector [9]. All point mutations were created using site-directed mutagenesis.

### Protein Expression and Purification

WT CaMKIIα kinase domain (residues 7-274) was co-expressed with λ phosphatase in *E. coli* BL21(DE3). Inactive constructs (D135N) and CaMKIIN2 *(Rattus norvegicus)* were expressed without λ phosphatase. The cells were induced at 18 °C with 1 mM isopropyl β-a-D-1-thiogalactopyranoside (IPTG) and grown overnight. Following to ∼16 hours incubation, cell pellets were resuspended in Buffer A (25 mM Tris, pH 8.5, 50 mM KCl, 40 mM imidazole, 10% glycerol) and commercially available protease inhibitors (0.5 mM benzamidine, 0.2 mM 4-(2-aminoethyl)benzenesulfonyl fluoride hydrochloride (AEBSF), 0.1 mg/mL trypsin inhibitor, 0.005 mM leupeptin, 1 µg/mL pepstatin), 1 µg/mL DNase/50 mM MgCl_2_ were added, then lysed. All following purification steps were performed using an ÄKTA pure chromatography system at 4 °C. Filtered cell lysate was loaded onto a 5 mL His Trap FF Ni Sepharose column (GE Research), and eluted with 50% Buffer B (25 mM Tris-HCl pH 8.5, 150 mM KCl, 1 M imidazole, 10% glycerol). The protein was desalted from excess imidazole using a HiPrep 26/10 Desalting column using Buffer C (25 mM Tris-HCl pH 8.5, 40 mM KCl, 40 mM imidazole, 2 mM tris(2-carboxyethyl)phosphine (TCEP), 10% glycerol). His SUMO tags were cleaved with Ulp1 protease overnight at 4 °C. Cleaved His SUMO tags were separated by a subtractive NiNTA step prior to an anion exchange step. Proteins were eluted from HiTrap Q-FF with a KCl gradient. Eluted proteins were concentrated and further purified in gel filtration buffer (25 mM Tris-HCl pH 8.0, 150 mM KCl, 1 mM TCEP, 10% glycerol) using Superdex 75 10/300 GL size exclusion column. Fractions (>95% purity) were pooled, concentrated, aliquoted and flash frozen in liquid nitrogen, and stored at -80 °C until needed.

We use rat full-length CaMKIIN2 protein in our ITC measurements, coupled-kinase assays and cellular immunoprecipitation experiments. The CaMKII-bound CaMKIIN co-crystal structure (PDB:3KL8) was obtained by using a peptide derived from rat CaMKIIN1. In our MD simulations, the full length human CaMKIIN1 amino acid sequence was used. Amino acid sequences of each construct are listed below:

CaMKIIN2 (*Rattus norvegicus*):

MSEILPYGEDKMGRFGADPEGSDLSFSCRLQDTNSFFAGNQAKRPPKLGQIGRAKRVVIEDDRIDDVLKGMGEKPPSGV

CaMKIIN1 peptide (*Rattus norvegicus*):

KRPPKLGQIGRSKRVVIA

CaMKIIN1 (*Homo sapiens*):

MSEVLPYGDEKLSPYGDGGDVGQIFSCRLQDTNNFFGAGQNKRPPKLGQIGRSKRVVIEDDRIDDVLKNMTDKAPPGV

### Peptide synthesis

Peptides used for co-crystallization were synthesized and amidated at the C-terminus (Genscript, RIKEN Research Resource Center). Peptide used for fluorescence polarization assays were synthesized with an additional N-terminal 5-FAM.

Full length peptide sequences are as follows:

Human GluA1 (residues 818-837): SKRMKGFCLIPQQSINEAIR [Numbering for GluA1 is for mature peptide, excluding 18 amino acid long signal peptide, following convention in the field.]

Human GluN2B (residues 1289-1310): KAQKKNRNKLRRQHSYDTFVDL

Human GluN2B(S1303D) (residues 1289-1310): KAQKKNRNKLRRQHDYDTFVDL

Human densin-180 (residues 797-818): SKSRSTSSHGRRPLIRQDRIVG

Mouse Tiam1 (residues 1541-1559): RTLDSHASRMTQLKKQAAL

Syntide-2: PLARTLSVAGLPGKK

Extended Syntide-2: GRRPLARTLSVAGLPGKK

Full length peptide sequence for fluorescence polarization assays are as follows:

Human GluN2B(S1303D) (residues 1289-1310): KAQKKNRNKLRRQHDYDTFVDL

### Crystallization and X-Ray Data Collection

Initial crystallization screening was done using the sitting vapor diffusion method at 4°C with commercially available screening kits. If needed, initial hits were optimized by the hanging vapor diffusion method. Final conditions for crystal formation and cryo solutions are listed below, all grown at 4°C. The peptide-to-protein ratio was kept at 3:1 throughout the co-crystallization attempts. ATP-bound crystal structures were obtained by adding 0.2 µL of reservoir solution containing 10 mM ATP/40 mM MgCl_2_ to crystal drops. For crystals that had methyl 6-O-(N-heptylcarbamoyl)-α-D-glucopyranoside (hecameg), a detergent in the crystallization condition (a detergent), the crystals were moved to a new drop without Hecameg, and containing 1 mM ATP/4 mM MgCl_2_. Diffraction data were collected from crystals flash-frozen in liquid nitrogen at a wavelength of 1.5418 Å using a Rigaku MicroMax-007 HF X-ray source, which was coupled to a Rigaku VariMax HF optic system (UMass Amherst). The X-ray data was collected at 100 K.

### Crystallization and cryo conditions are listed below

6X5G: 0.1 M Bicine pH 9, 10% PEG 20K, 2% 1,4-Dioxane (Cryoprotectant: 20% glycerol)

6X5Q: 0.1 M Tris pH 8, 28% PEG 4K (Cryoprotectant: 20% glycerol)

7UIQ: 0.1 M HEPES pH 7, 16% PEG 6K, 0.1% Triton X-114 (Cryoprotectant: 20% ethylene glycol)

7UJQ: 0.1 M Bis-tris methane pH 6.2, 0.1 M ammonium sulfate, 25% PEG 3350, 19mM Hecameg (Cryoprotectant: 20% glycerol)

7UJP: 0.1 M Bis-tris methane pH 6.2, 0.1 M ammonium sulfate, 25% PEG 3350, 19mM Hecameg (Cryoprotectant: 20% glycerol)

7UJR: 0.1 M Bis-tris propane pH 6.2, 0.1 M ammonium sulfate, 20% PEG 3350, 5% ethanol (Cryoprotectant: 20% ethylene glycol)

7UJS: 0.1 M Bis-tris propane pH 6.6, 0.1 M ammonium sulfate, 20% PEG 3350 (Cryoprotectant: 20% ethylene glycol)

7UIR: 0.1 M HEPES pH 7, 16% PEG 6K, 0.1% Triton X-114 (Cryoprotectant: 20% ethylene glycol)

7UJT: 0.1 M Bis-tris propane pH 6.5, 0.2 M Sodium chloride, 25% PEG 3350 (Cryoprotectant: 20% glycerol)

7KL0: 0.1 M Bis-tris methane pH 6.5, 0.1 M Ammonium sulfate, 20% PEG 6000, 19mM Hecameg (Cryoprotectant: 20% ethylene glycol)

7KL1: 0.1 M Bis-tris methane pH 6.5, 0.1 M Ammonium sulfate, 20% PEG 6000, 19mM Hecameg (Cryoprotectant: 20% ethylene glycol)

7UIS: 0.1 M Bis-tris propane pH 6.5, 0.2 M Sodium chloride, 25% PEG 3350 (Cryoprotectant: 20% glycerol)

### Data Processing and Structure Determination

Data sets were integrated, merged, and scaled using HKL-2000. The structures were solved by molecular replacement (MR) with Phaser using the coordinates of CaMKIIα kinase domain (PDB ID: 6VZK, 100% amino acid sequence identity) as a search model [48]. Peptides were built into electron density using Coot and refinement was performed with REFMAC5 [49, 50].

### Isothermal titration calorimetry

ITC data were obtained using a MicroCal PEAQ-ITC automated calorimeter (Malvern Panalytical, Westborough, MA). Before each titration, interaction partners were dissolved in gel filtration buffer and final buffer conditions of CaMKII kinase domain and interaction partners were matched. GluN2B peptides and CaMKIIN protein contain a single tyrosine residue, which enabled accurate concentration determination. The other peptides lack aromatic residues, and these required crude weight to estimate concentration. Titrations were performed with different peptides as titrant into the cell containing D135N or other mutant CaMKII kinase domains (concentrations are indicated in Table 3). All titrations were performed using the standard 19 injection method, modified to run at 20 °C. The standard 19 injection method in the PEAQ ITC automated control software (v1.40) is one injection of 0.4 µL followed by 18 2 µL injections with a spacing between injections of 150s, a stir speed of 750 rpm, and the reference power set to 10. Data were analyzed using PEAQ ITC analysis software (v1.40) using the one-site fitting model, and to eliminate integration of some baseline noise, some integration markers were set manually to obtain better fit.

### Fluorescence polarization assay

The competition assay was conducted at 1 µM of CaMKII kinase domain (25 mM Tris pH 7.5, 150 mM KCl, 1 mM TCEP, 10% glycerol) with 60 nM fluorescein-labeled GluN2B peptide (dissolved in 25 mM Tris pH 7.5, 150 mM KCl, 0.02% Tween, 0.02% Triton) and then titrated increasing concentrations of unlabeled peptide (25 mM Tris pH 7.5, 150 mM KCl, 1 mM TCEP, 10% glycerol) at varying concentrations in Corning low volume 384-well black flat bottom plates. The final volume in each well was 20 µL. The fluorescence polarization was measured using a Synergy H1 hybrid plate reader (Biotek) with a filter of 485/20 nm excitation and 528/20 nm emission. Data were fit using a one-site binding equation in GraphPad PRISM version 6.0.

### Differential scanning calorimetry

All protein samples were diluted to 0.5 mg/mL in DSC buffer (25 mM Tris pH 8,150 mM KCl, 1 mM TCEP, 10% glycerol). DSC measurements were performed on a MicroCal Automated PEAQ-DSC instrument (Malvern Panalytical, Westborough, MA). Unless otherwise indicated, after a 5 min pre-scan equilibration step, samples were scanned from 10-120°C at a scan rate of 90°C/hr with no feedback. Data were analyzed using MicroCal PEAQ-DSC software, and baseline-subtracted data were fit to a non-two-state fitting model to obtain apparent T_m_ values.

### Molecular Dynamics simulations

Trajectories were run for 1.2 μs - 3.7 μs aggregate time for each complex. All simulations are available for download from Mendeley: DOI: 10.17632/9m3bg3h5rs.1. Starting conformations were taken from co-crystal structures of CaMKIIα kinase domain in complex with peptides derived from GluN2B (PDB:7UJP) and Tiam1 (PDB:7UIR). The sequences used for interaction partners in the simulations are as follows: Tiam1 (1513-1581, adding 28 N-terminal residues and 22 C-terminal residues) as well as GluN2B (1263-1328, adding 32 N-terminal residues and 20 C-terminal residues), and the full-length CaMKIIN protein (adding 41 N-terminal residues and 22 C-terminal residues). For CaMKIIN, Coot [50] was used to superimpose the structure of a peptide derived from rat CaMKIIN in complex with the kinase domain from *C. elegans* CaMKII (PDB:3KL8) with the structure of GluN2B in complex with the kinase domain of CaMKIIα bound to a Mg^2+^ ion and an ATP molecule (PDB:7UJP). The CaMKIIN peptide and the CaMKIIα kinase domain were then merged into a single structure. Crystallographically unresolved atoms were added, peptides were extended in both N-terminal and C-terminal directions using PDBFixer, part of the OpenMM package [51].

All simulations were performed with Gromacs 2020.2 [52, 53] and the CHARMM36m force field [53]. Protein complexes were solvated with TIP3P water [54] in dodecahedral boxes that extended 1 nm past the protein in all dimensions. Na^+^ ions were added in sufficient quantity to charge neutralize each system. Systems were energy-minimized by steepest descent for 50,000 steps using a 0.01 nm step size until the maximum force exerted on any atom fell below 1000 kJ/mol/nm. Solvent was equilibrated for 1 ns at constant temperature and volume (NVT ensemble) and another 2 ns at constant temperature and pressure (NPT ensemble) with a positional restraint applied to protein heavy atoms. An additional 10 ns of unrestrained dynamics were simulated prior to collection of production data in order to allow the polypeptide chains to relax from their crystallographic conformations, and allow the simulated systems to equilibrate fully. Periodic boundary conditions were used in all dimensions, bonds were constrained with the LINCS algorithm [55], virtual sites (v-sites) were used to remove the fastest degrees of freedom to facilitate a 4 fs time step [56], particle mesh Ewald (PME) was used to treat electrostatic interactions [57], the v-rescale thermostat [58] with a 0.1 ps coupling time constant was used to maintain the temperature at 300 K, and a cut-off distance of 1.2 nm for neighbor list, Coulomb interactions, and Van der Waals interactions was used throughout system preparation and production simulations. The Parrinello–Rahman barostat [59] with a 2 ps coupling time constant was used to maintain a pressure of 1 bar during NPT equilibration and production simulations. V-site parameters for ATP were determined using MkVsites [60]. GPU-accelerated simulations were performed on two hardware architectures: 1) 2x Intel Xeon Silver 4110 cpu (8 Cores), 2x nVidia Tesla V100 gpu; 2) 2x Intel Xeon Gold 6140 cpu (18 Cores), 2x nVidia Tesla V100 gpu. In order to obtain comparable aggregate lengths of production simulations for the three simulated systems, two independent simulations of the kinase:CaMKIIN complex were carried out.

To calculate RMSD and RMSF during the simulations, trajectories were aligned about the Cα of the kinase domain. RMSD was calculated with VMD [61]. RMSF was calculated with ProDy [62] as interfaced with VMD. RMSD and RMSF of the peptide were calculated with the kinases aligned to determine how the residues of the peptide fluctuated with respect to the kinase.

One production simulation was performed for each of the wild type kinase:GluN2B and wild type kinase:Tiam1 complexes, producing trajectories of 2.19 μs and 2.05 μs, respectively. Two independent simulations were performed for the wild type kinase:CaMKIIN. The CaMKIIN trajectories were 0.46 μs and 1.84 μs in length, for an aggregate of 2.3 μs. The GluN2B trajectories were 0.68 μs and 0.57 μs in length, for an aggregate of 1.25 μs. Trajectories were analyzed using MDTraj [63]. Hydrophobic contact distance between two residues was defined as the shortest distance between side chain heavy atoms. Pairwise RMSD between frames for specific hydrophobic interactions (kinase residues F98, I101, V102, I205, and peptide residue in -5 position adjusted from sequence alignment; kinase residue W214, Ile at –5, Pro at –10 in CaMKIIN peptide, see Fig 2B) was calculated based on side chain heavy atoms for the residues involved in the interaction: 5 residues for the first interaction, 3 residues for the second.

### Coupled kinase assays

Kinase activity was monitored using a Synergy H1 microplate reader (Biotek) as previously described [22, 64]. The assay was conducted in 50 mM Tris, 150 mM KCl, 10 mM MgCl_2_, 2 mM ATP, 1 mM Phosphoenolpyruvate (Alfa Aesar), 0.2 mM Nicotinamide adenine dinucleotide (Sigma), 10 units/mL Pyruvate kinase (Sigma), 30 units/mL Lactate dehydrogenase (Millipore Sigma), varying concentrations of Syntide-2 (Lifetein). The final CaMKII kinase domain concentration was 5 nM. 500 nM CaMKIIN protein or 8.14 μM NMDAR(S1303D) peptide was preincubated with CaMKII kinase domain before adding to the reaction mix of corresponding experiments. The reactions were started by the addition of ATP to the reaction mix and the absorbance was measured at 340 nM at 30°C at 10 and 15 sec intervals for 10 minutes. The rate was obtained by calculating the maximum observed slope of each reaction. Data were fit using the Michaelis-Menten equation in GraphPad PRISM version 6.01.

### Immunoprecipitation

HEK293T cell lysates were prepared in lysis buffer (50 mM Tris-HCl pH 7.5, 150 mM NaCl, 1% Triton X-100, 10% glycerol, 1 mM Na_3_VO_4_, 10 mM NaF, 1 mM β-glycerophosphate, 1 x phosphatase inhibitor cocktail (Nacalai, Kyoto, Japan), 1x cOmplete tablet (Roche, Basel, Switzerland) and centrifuged at 14,000 rpm for 10 min at 4°C. The supernatant was subjected to immunoprecipitation using 20 μL of the anti-Flag antibody-beads (Sigma) for 2-4 hours at 4°C.

Beads were washed with 1 mL of lysis buffer for three times. Bound proteins were eluted with SDS-PAGE sample buffer and subjected to western blotting.

## Data availability

Atomic structures have been deposited in the PDB, listed in the table below.

Table of PDB codes and descriptions.

|  |  |
| --- | --- |
| 6X5G | D135N kinase in complex with densin-180 |
| 6X5Q | D135N kinase in complex with GluA1 |
| 7UIQ | D135N kinase in complex with Tiam1 |
| 7UIR | D135N kinase in complex with Tiam1 and ATP |
| 7UJQ | D135N kinase in complex with GluN2B and Hecameg |
| 7UJP | D135N kinase in complex with GluN2B and ATP |
| 7UJR | WT kinase in complex with GluN2B |
| 7UJS | WT kinase in complex with GluN2B and ADP |
| 7KL0 | D135N kinase in complex with phosphomimetic GluN2B and Hecameg |
| 7KL1 | D135N kinase in complex with phosphomimetic GluN2B and ATP |
| 7UIS | D135N kinase in complex with phosphomimetic GluN2B |
| 7UJT | D135N kinase in complex with phosphomimetic GluN2B and ATP |

## ACKNOWLEDGMENTS

We thank Roger Colbran, Howard Schulman, Eric Strieter and Peter Chien for helpful discussions. We thank Josh Pajak for help with MD simulations. This work was supported by funds from NIGMS R01123157 (M.S.), SPIRITS 2019 of Kyoto University, Grant-in-Aid for Scientific Research JP18H04733, and JP18H05434 from the MEXT, CREST JPMJCR20E4 from JST, Japan and HFSP Research Grant (RGP0020/2019) (Y.H.).

## Author Contributions

C.Ö. solved all crystal structures and performed all FP measurements and kinetic assays. T.M. and T.S. performed immunoprecipitation experiments. N.S., E.A., C.G., E.L., J.F. and E.A.E. also assisted with experiments under supervision of B.A.K., S.C.G., Y.H. and M.M.S. R.S. performed molecular dynamics. C.Ö., R.S., Y.H., B.A.K., and M.M.S. wrote and edited the manuscript.

## SUPPLEMENTAL FIGURES

**Figure S1.**
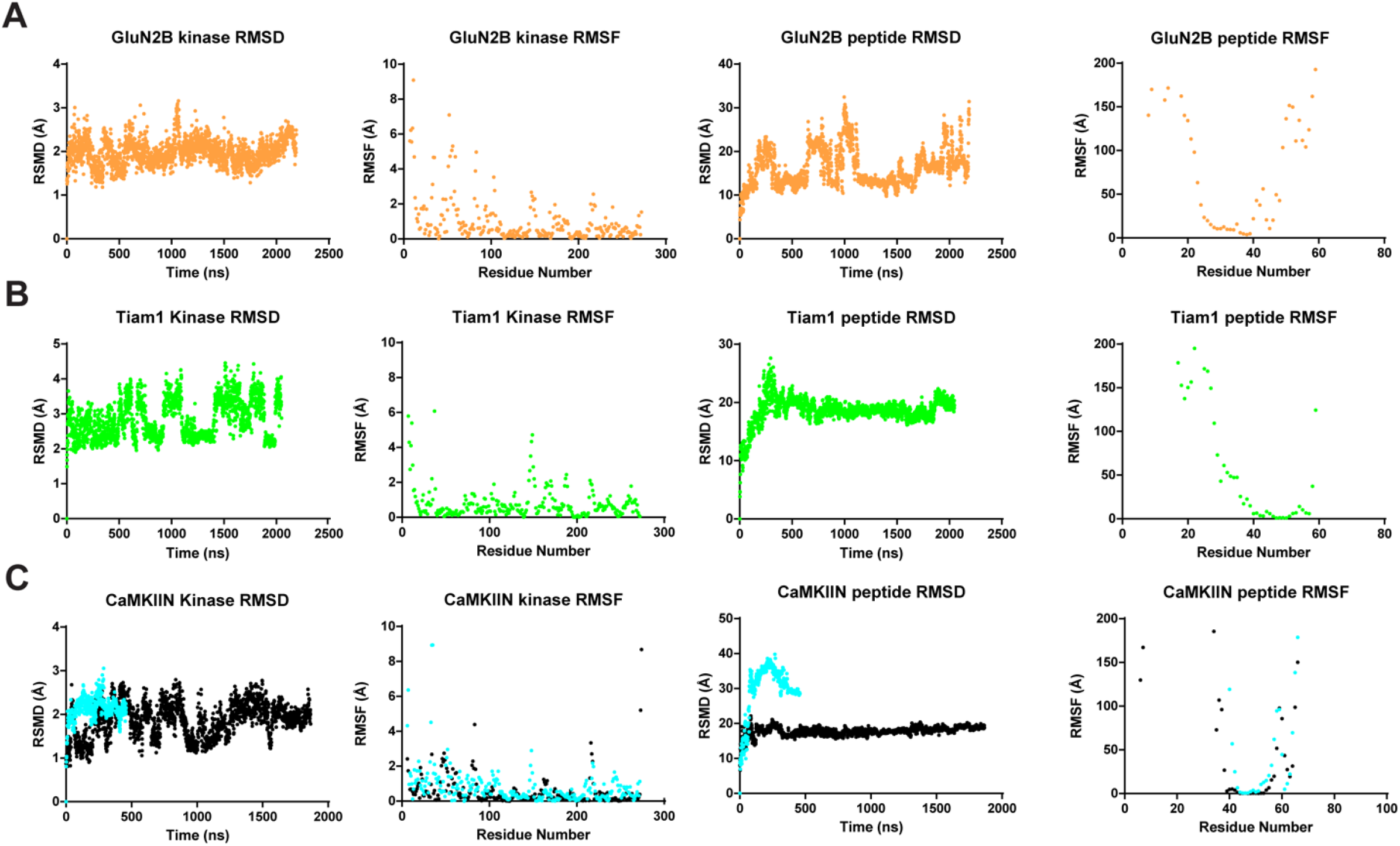
RMSD and RMSF calculations from MD trajectories. RMSD and RMSF were calculated for both the kinase domain and the peptide for each of the trajectories. For CaMKIIN, the two trajectories are overlaid (cyan:0.46 μs and black:1.84 μs). For the kinase domains, RMSD and RMSF values are largely ∼3 Å or less, with higher values at the termini, indicating high stability. For the peptides, RMSD values are higher which is not surprising considering this is a largely flexible polypeptide. Peptide RMSF plots are shown up to 200 Å. In the bound regions (residues ∼23-40 in GluN2B, ∼40-60 in Tiam1 and CaMKIIN) the RMSF values are quite low (<4 Å) indicating high stability. Whereas there is high mobility at the termini.

**Figure S2.**
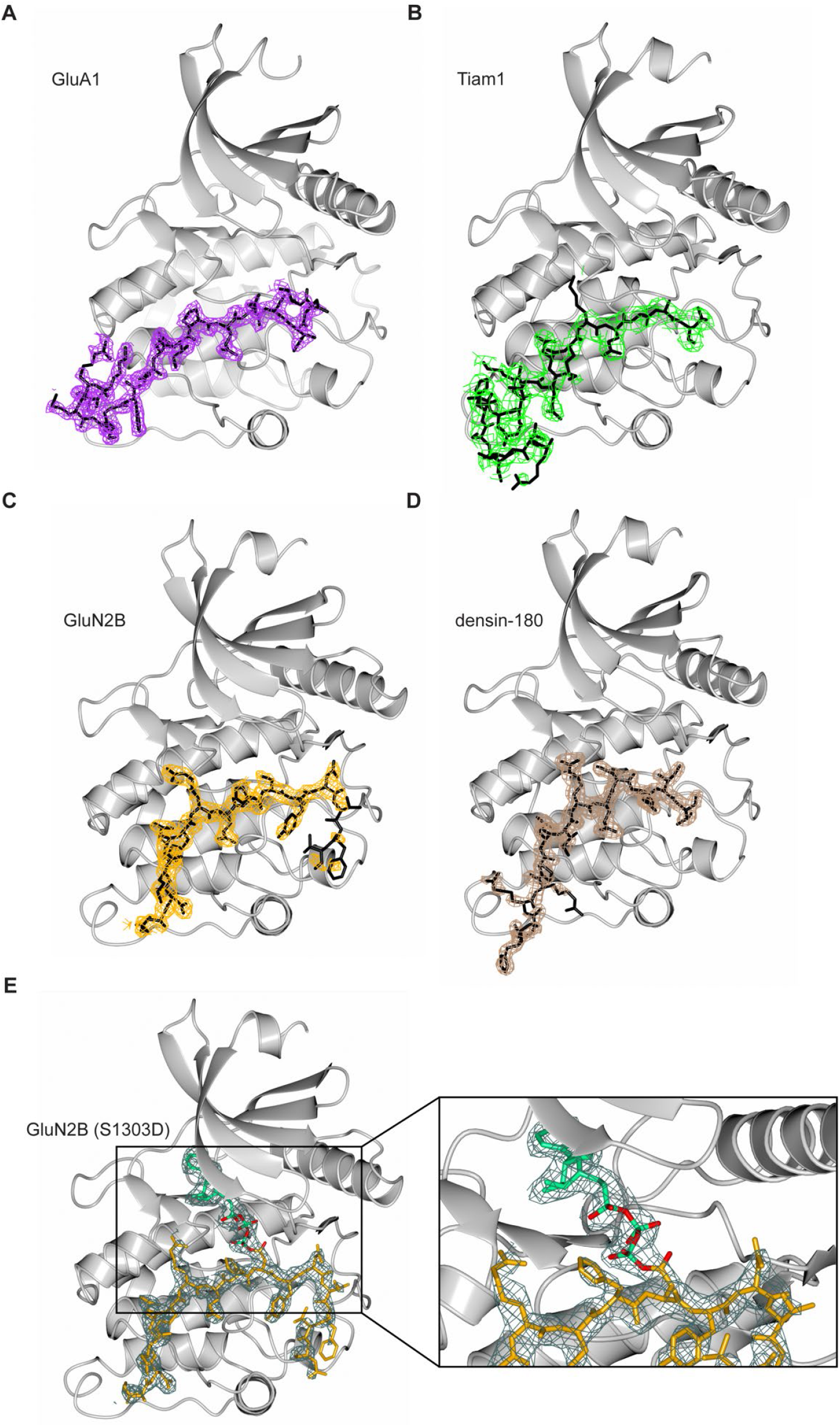
Structures of CaMKII kinase domain bound to peptide binding partners. Electron-density omit maps are shown as mesh for all peptide binding partners at σ=1. CaMKII kinase domain shown in light gray (A) GluA1 (PDB:6X5Q) in purple, (B) Tiam1(PDB:7UIQ) in green, (C) GluN2B (PDB:7UJQ) in orange, (D) densin-180 (PDB:6X5G) in brown, and (E) GluN2B (S1303D) in presence of ATP (PDB:7KL1). For (E), the peptide is shown in orange and omit map is shown in dark gray, ATP is colored green. D1303 sidechain is covalently linked to ATP molecule and is shown in red together with the phosphate oxygens.

**Figure S3.**
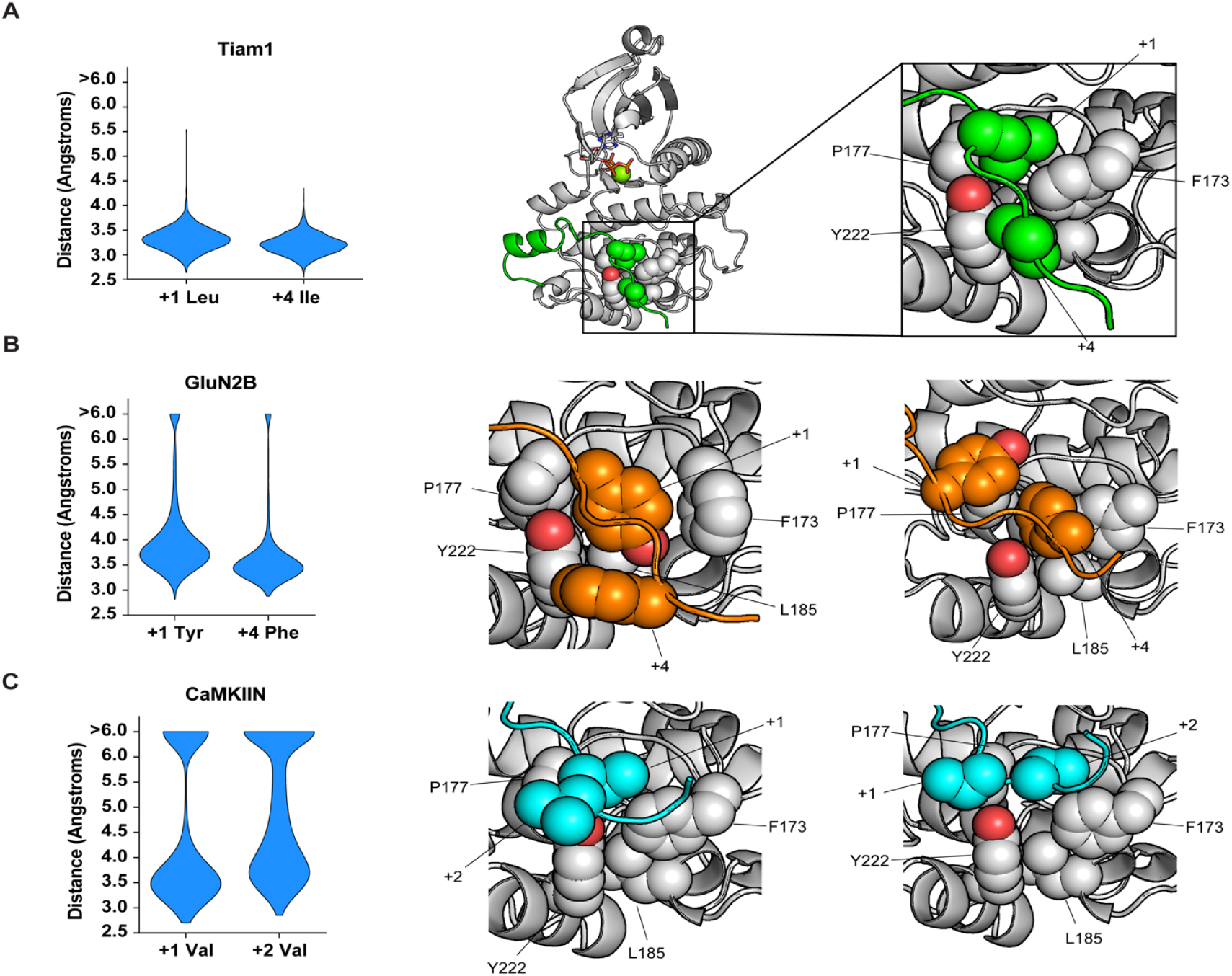
MD simulations highlight a hydrophobic interaction at the C-terminal end of peptides. (A) Persistence of Tiam1 interaction with the F173, P177, L185, Y222 hydrophobic patch in MD simulations. Distance distributions for +1 leucine and +4 isoleucine of Tiam1 to patch (left), representative structure (middle). Zoomed view is labeled for clarity. (B) Persistence of GluN2B interaction with hydrophobic patch. Distance distributions for +1 tyrosine and +4 phenylalanine of GluN2B and two zoomed-in views of the interaction from different orientations. (B) Persistence of CaMKIIN1 interaction with hydrophobic patch from different orientations. Distance distributions for +1 and +2 valine residues of CaMKIIN and two zoomed-in views of the interaction from different orientations.

**Figure S4.**
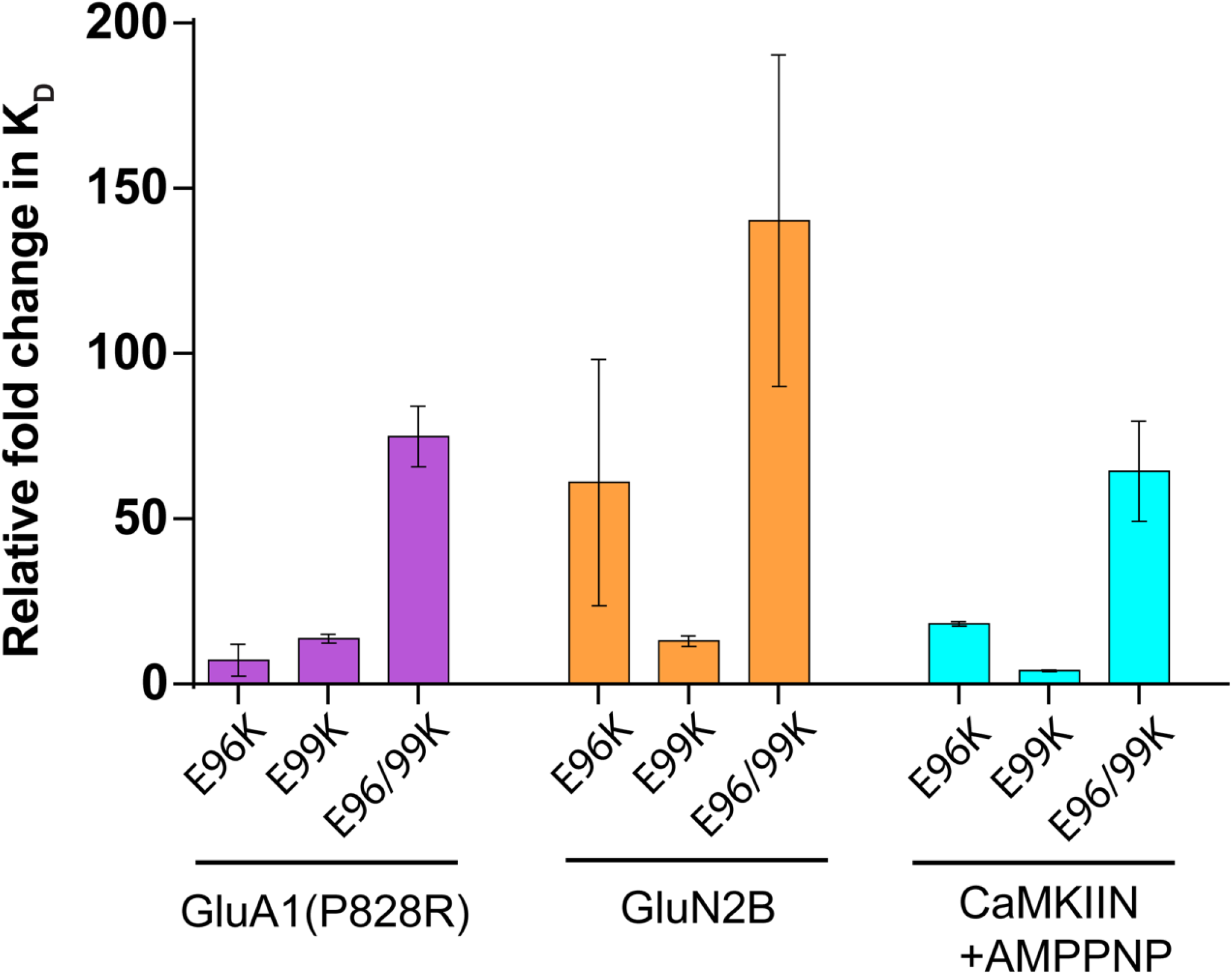
Effect of charge reversal mutations at the -3 position. K_d_ values were extracted from ITC data and relative fold changes were calculated by dividing the observed K_d_ from the mutant by the D135N kinase domain. Standard deviations are shown as error bars for duplicate measurements.

**Figure S5.**
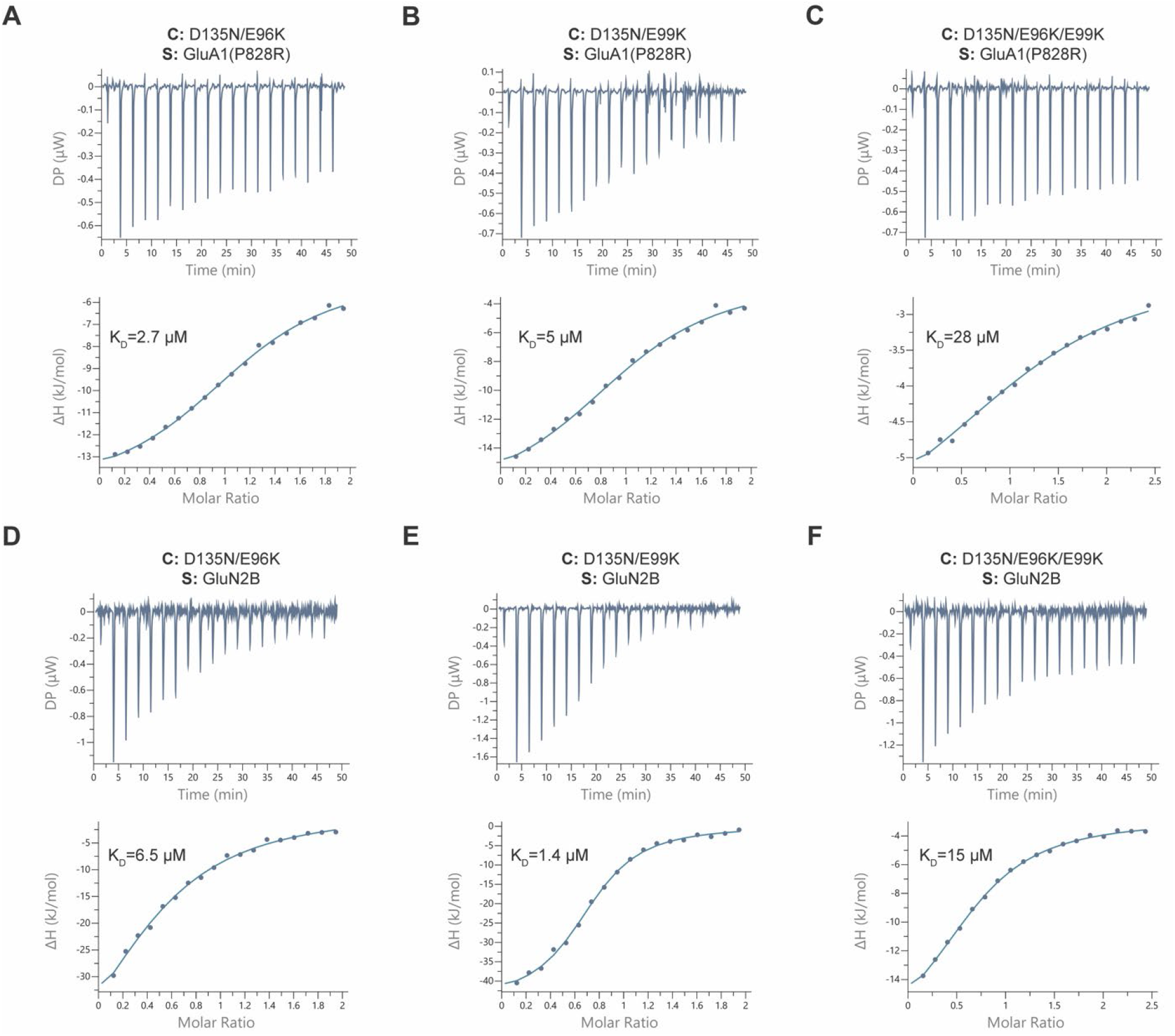
Effect of E96K and E99K mutations on binding affinity. ITC measurements of D135N kinase domain mutants and interaction partners. All measurements were performed in duplicate, here one representative dataset is shown. (A) E96K and GluA1 (P828R), (B) E99K and GluA1 (P828R), (C) E96K/E99K and GluA1 (P828R), (D) E96K and GluN2B, (E) E99K and GluN2B, and (F) E96K/E99K and GluN2B. The mean K_d_ value from two independent measurements is labeled in the figure.

**Figure S6.**
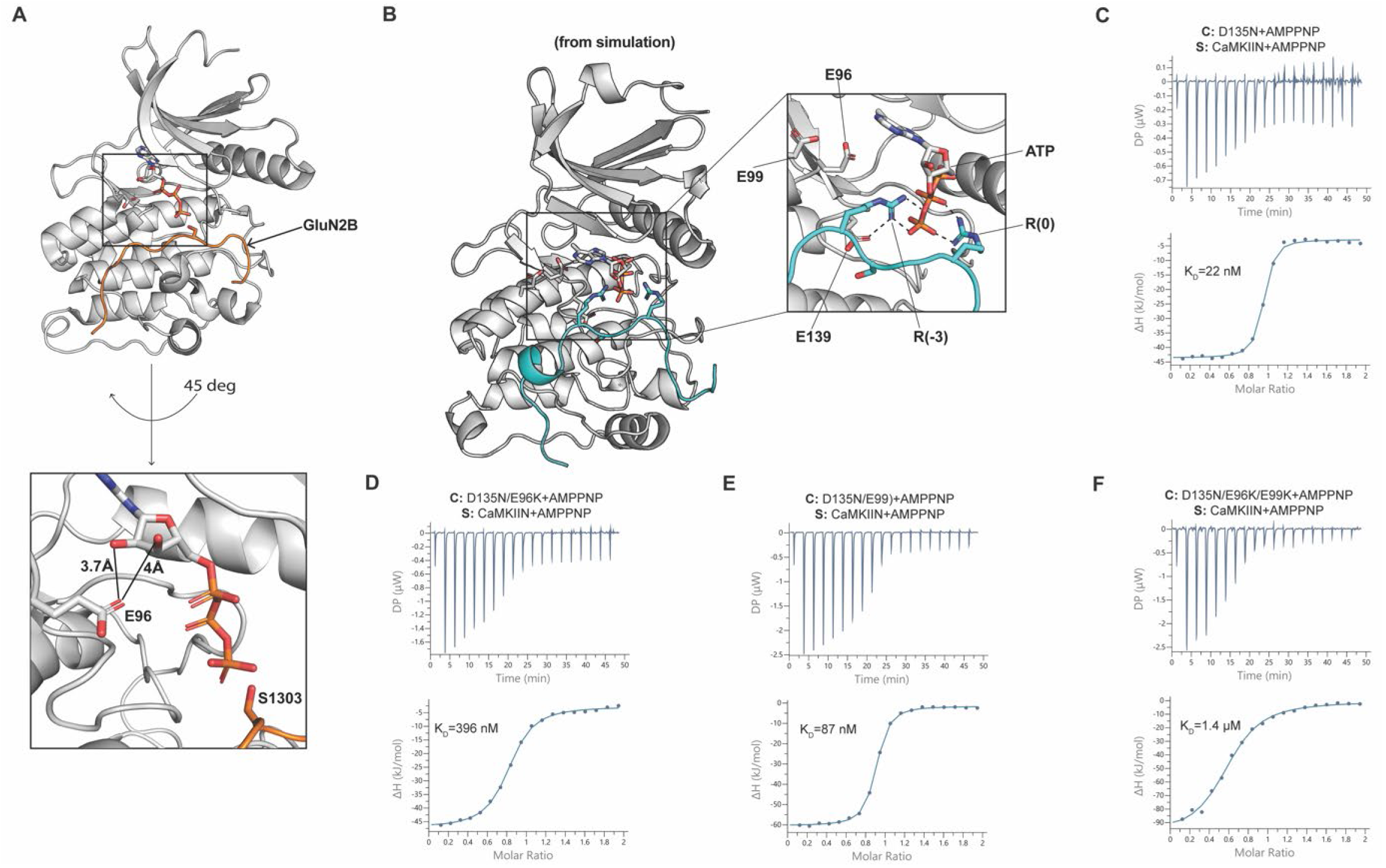
CaMKIIN has unique interactions with ATP. (A) Co-crystal structure of CaMKII kinase domain and GluN2B in complex with ATP (PDB 7UJP). Bottom image highlights the interaction between E96 sidechain and ribose hydroxyl groups of ATP. The gamma phosphate group faces S1303. (B) Snapshot from an MD trajectory with CaMKIIN peptide and ATP bound. The arginine at the -3 position of CaMKIIN mostly interacts with E139, minimally with E96, and not at all with E99. All interactions depicted with dashed lines are below 3 Å. This conformational change precludes the ATP adenosine from forming ionic interactions with the backbones of D90 and V92, as observed in the crystal structure. As a result, the ATP adenosine group exits the binding pocket in the CaMKIIN trajectories. ITC measurements of CaMKIIN in the presence of AMPPNP are shown in C-F. Contents of the cell (C) and syringe (S) used in the ITC measurements are listed in the figure panels. The mean K_d_ value from two independent measurements is labeled in the figure.

**Figure S7.**
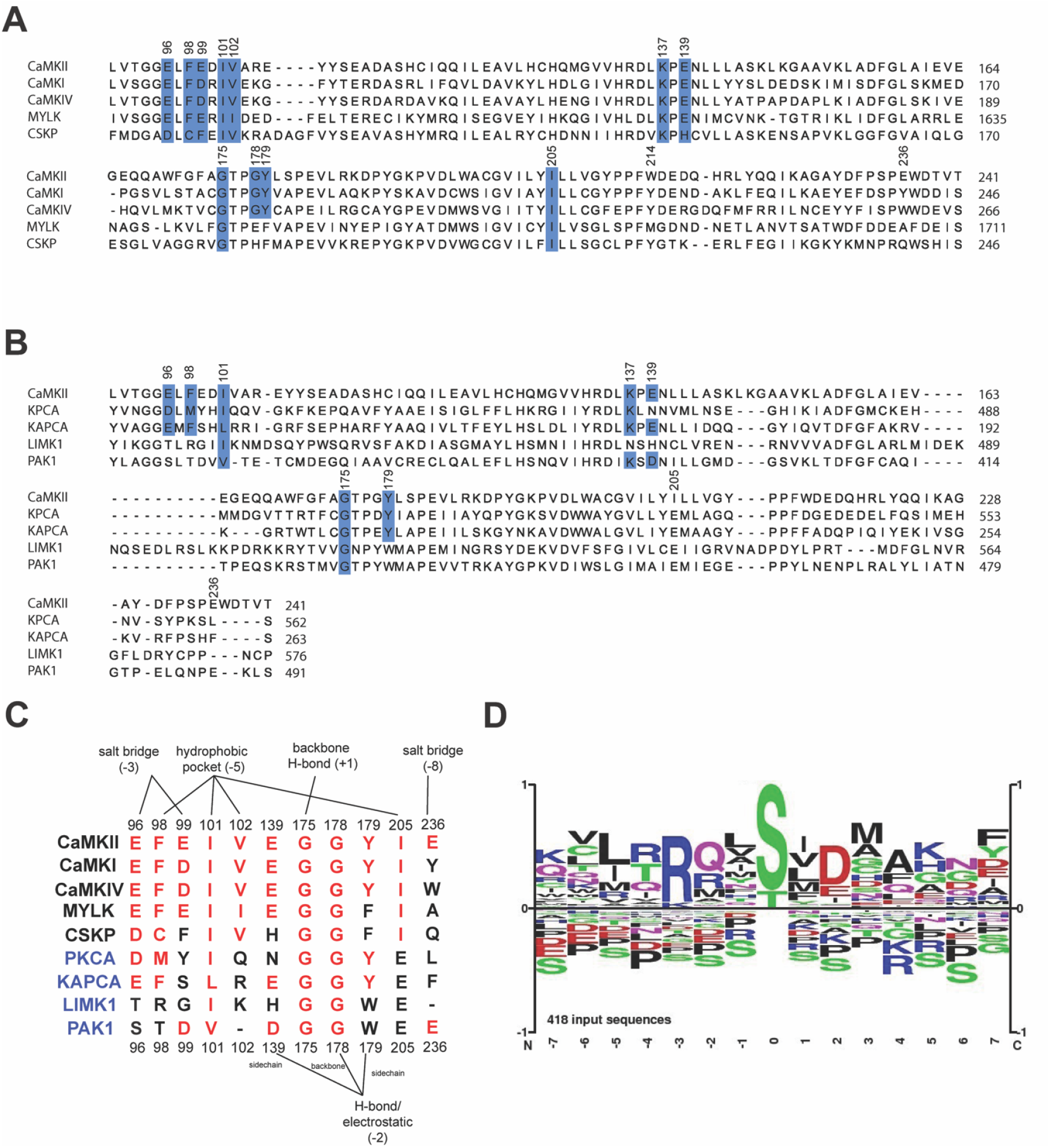
Multi-sequence alignment of kinase domains. Alignment of CaMKII from residue 91-241 (according to CaMKIIα numbering) is shown (A) across Ca^2+^/CaM (CaMK) family and (B) across different kinase families. Conserved residues are highlighted blue. (C) Alignment of key residues involved in binding substrates across kinases from the Ca^2+^/CaM family (black text) and four kinases from different families (blue text). PKCA and KAPCA (cAMP dependent kinase) belong to the AGC Ser/Thr family. LimK1 and Pak1 belong to TKL and STE Ser/Thr family. (D) The sequence logo for all previously identified substrates of CaMKII from the PhosphoSitePlus database with the phosphorylation site at the 0 position.

**Figure S8.**
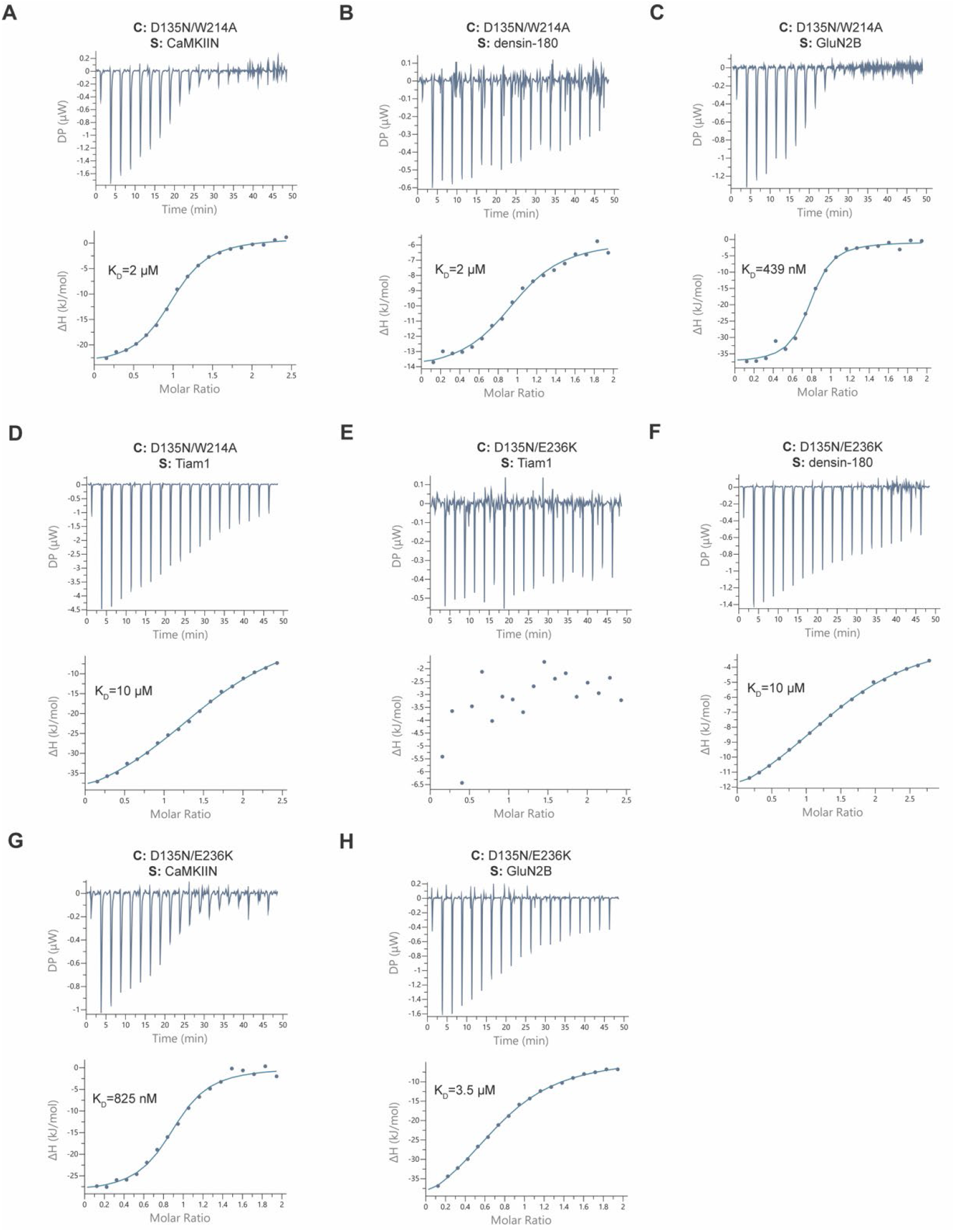
Effect of W214A and E236K mutations on binding affinity. ITC measurements between D135N kinase domain mutants and interaction partners. Contents of the cell (C) and syringe (S) used in the ITC measurements are listed in the figure panels. The mean K_d_ value from two independent measurements is labeled in the figure.

**Figure S9.**
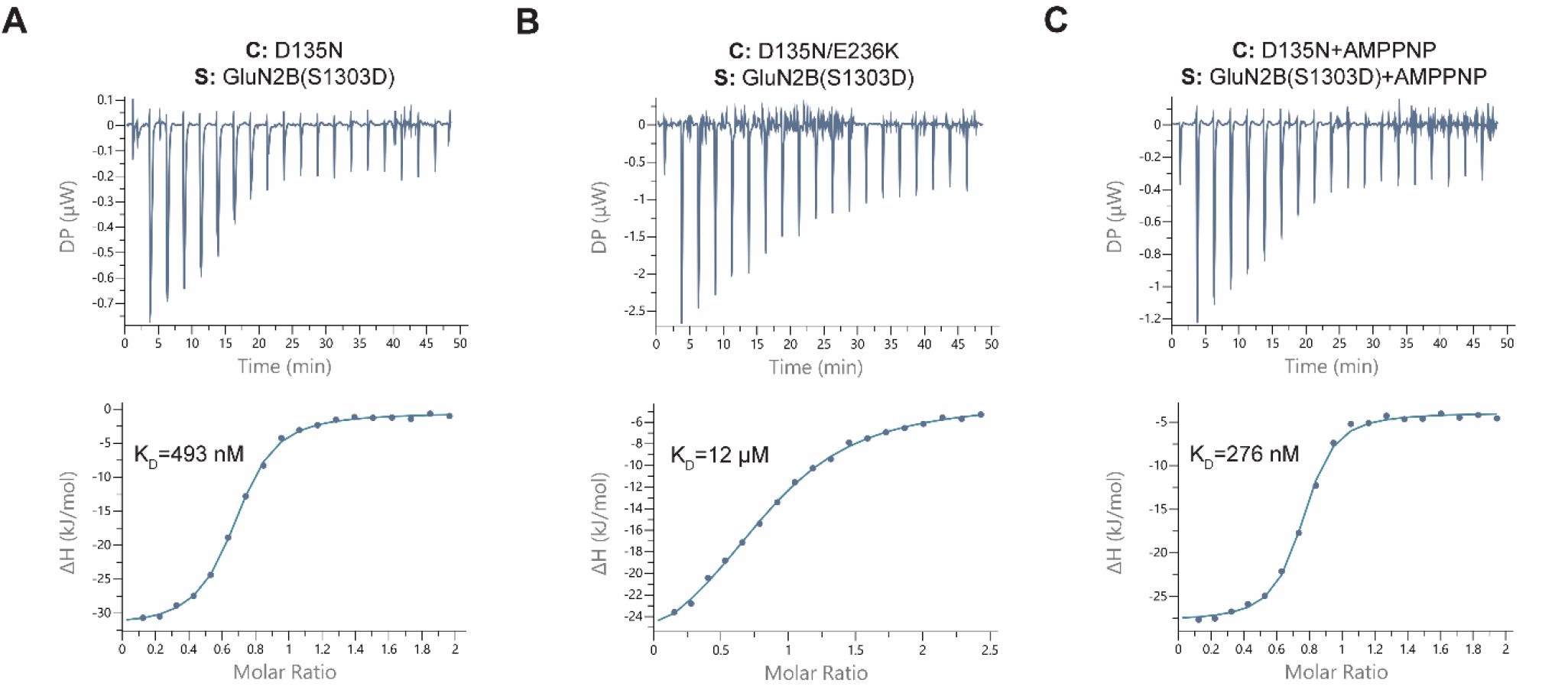
Effect of the phosphomimetic mutation on GluN2B binding affinity. ITC measurements between D135N kinase domain mutants and interaction partners. Contents of the cell (C) and syringe (S) used in the ITC measurements are listed in the figure panels. The mean K_d_ value from two independent measurements is labeled in the figure.

**Figure S10.**
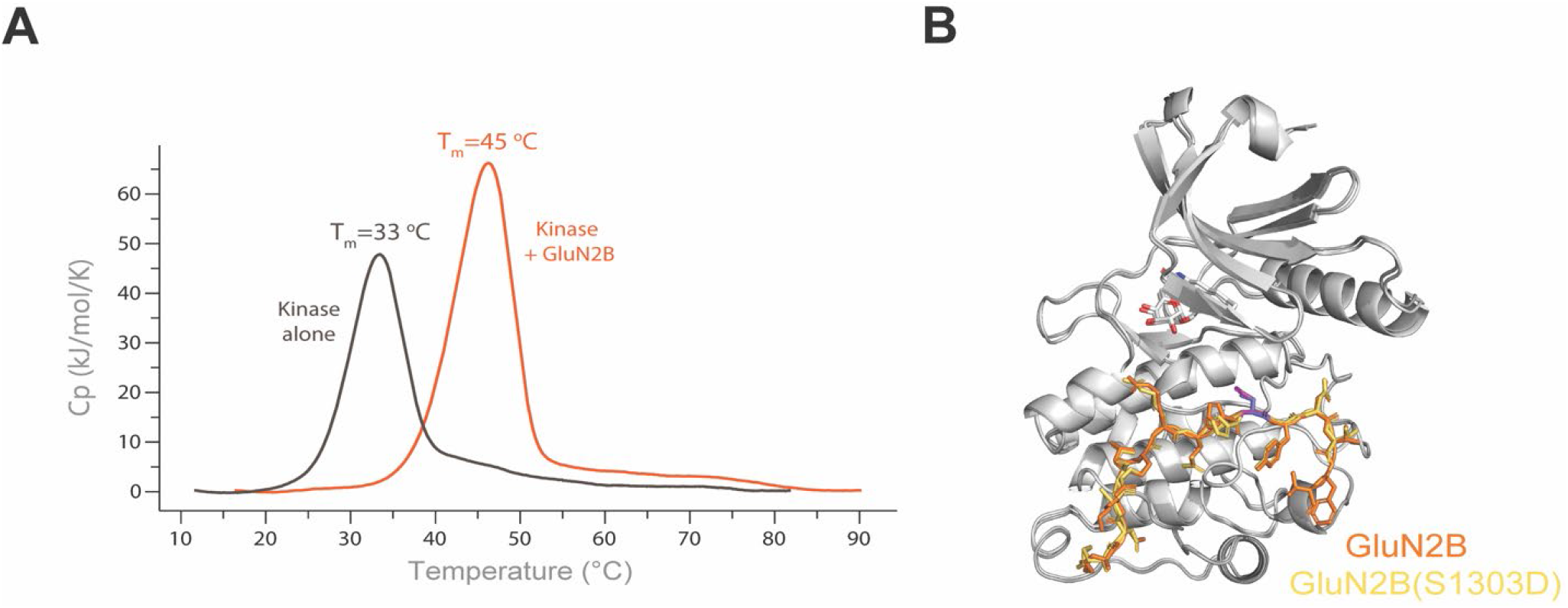
Alignment of WT and S1303D GluN2B bound to the kinase domain and DSC for GluN2B binding. (A) Differential Scanning Calorimetry data from CaMKII kinase domain alone (brown) and CaMKII kinase domain bound to GluN2B peptide (orange). (B) Overlay of WT and S1303D GluN2B bound kinase domain structures with hecameg bound (detergent present in crystallization condition) (PDB: 7UJQ and 7KL0). The residue at 1303 is highlighted in blue (serine) and magenta (aspartate).

**Figure S11.**
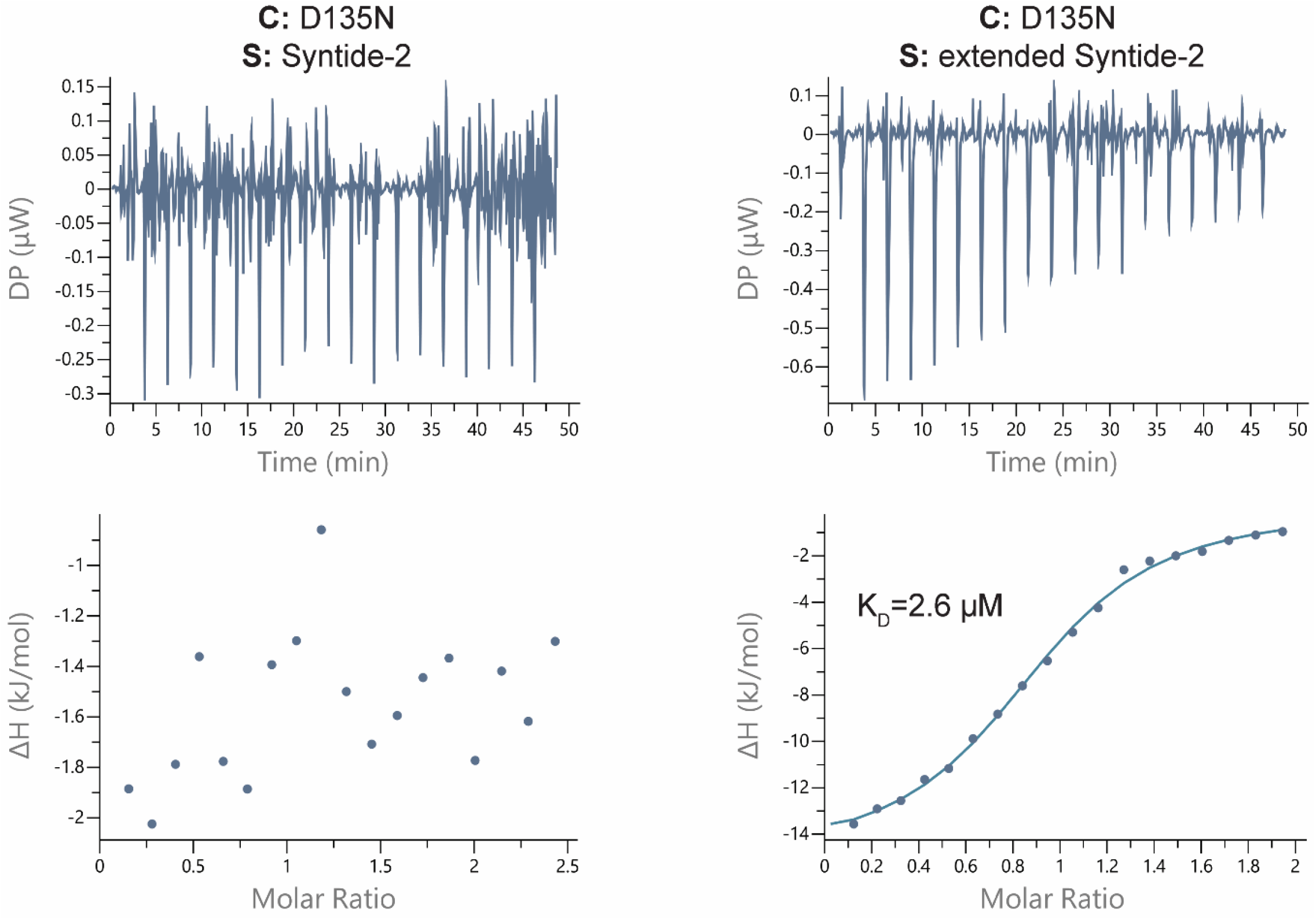
Far salt bridge interaction facilitates low micromolar affinity. Comparison of Syntide-2 to extended Syntide-2. Contents of the cell (C) and syringe (S) used in the ITC measurements are listed in the figure panels. The mean K_d_ value from two independent measurements is labeled.

